# Correlated evolution of beak and braincase morphology is present only in select bird clades

**DOI:** 10.1101/2023.11.19.567761

**Authors:** Xiaoni Xu, Rossy Natale

**Affiliations:** University of Chicago, Department of Organismal Biology and Anatomy, Chicago, Illinois 60637

**Keywords:** geometric morphometrics, Charadriiformes, macroevolution, modularity, integration

## Abstract

Complex morphological structures, such as skulls or limbs, are often composed of multiple morphological components (e.g. bones, sets of bones) that may evolve in a covaried manner with one another. Previous research has reached differing conclusions on the number of semi-independent units, or modules, that exist in the evolution of structures and on the strength of the covariation, or integration, between these hypothesized modules. We focus on the avian skull as an example of a complex morphological structure for which highly variable conclusions have been reached in the numerous studies analyzing support for a range of simple to complex modularity hypotheses. We hypothesized that past discrepancies may stem from both the differing densities of data used to analyze support for modularity hypotheses and the differing taxonomic levels of study. To test these hypotheses, we applied a comparative method to 3D geometric morphometric data collected from the skulls of a diverse order of birds (the Charadriiformes) to test support for 11 distinct hypotheses of modular skull evolution. Across all Charadriiformes, our analyses suggested that charadriiform skull evolution has been characterized by the semi-independent, but still correlated, evolution of the beak from the rest of the skull. When we adjusted the density of our morphometric data, this result held, but the strength of the signal varied substantially. Additionally, when we analyzed subgroups within the order in isolation, we found support for distinct hypotheses between subgroups. Taken together, these results suggest that differences in the methodology of past work (i.e. statistical method and data density) as well as clade-specific dynamics may be the reasons past studies have reached varying conclusions.

## 2 Introduction

The patterns defining the morphological evolution of complex or diverse structures have long been of interest in evolutionary biology. In particular, how the correlated evolution of different anatomical units may relate to the phenotypic space available to a species or to the rates of diversification in morphological structures are common questions (e.g. Navalón et al. 2020; Felice et al. 2018) in regards to morphological evolution. This idea is encapsulated by the term integration which, in the context of evolutionary morphology, describes the correlated evolution of traits within clades (Zelditch and Goswami, 2021). Differences in the degree of integration within and between different morphological features generate evolutionary modularity: the semi-independent evolution of sets of traits (Klingenberg 2008). A greater degree of modularity represents stronger evolutionary covariation between individual components within a module relative to the associations between modules (Adams and Collyer, 2019). Modularity is often examined in the context of other factors, such as diversification rates to provide insight into the mechanisms governing the diversification of complex morphological structures. For example, Larouche et al. (2018) found support for modular evolution across the bodies of ray-finned fishes and, by analyzing body shape evolution in a phylogenetic context, were able to make predictions about how increased modularity may facilitate rapid morphological diversification.

The vertebrate skull has been a focus of research on evolutionary modularity due to its heterogeneous developmental origins (Couly GF, 1993) and its multiple roles in biomechanical (e.g. feeding on different food items; Vidal-Garćıa and Scott Keogh 2017) and sensory systems (e.g. vision; Heesy 2008). Morphometric data have been used to test support for different modularity hypotheses across the skulls of diverse clades, including amphibians (Bardua et al., 2019; Bon et al., 2020), mammals (Adams and Collyer, 2019; Goswami and Finarelli, 2016), and birds (Klingenberg and Marugán-Lobón, 2013). These studies have found support for modular hypotheses of varying complexities, with studies often recovering the strongest support for hypotheses with a large number of semi-independent skull modules. For example, Watanabe et al. (2019) found support for a hypothesis in which the skulls of lizards and snakes evolved as 10 and 9 semi-independent modules, respectively, while analyses by Bardua et al. (2020) suggest an even more complex 13 module patterns in frog skull evolution. These findings suggest that the patterns of modular evolution vary across clades of vertebrates, but may often involve very finely subdivided parts of the skull evolving semi-independently from one another.

Underlying these studies of modularity are often interests in understanding how the semi-independent evolution of anatomical regions may relate to the generation of ecologically important morphological diversity. It has been hypothesized that the pronounced variation in avian beak shape (Cooney et al., 2017) may have been facilitated by its semi-independent evolution from the rest of the avian skull. Application of partial least squares (PLS) analysis, as well as calculations of RV coefficients on 2-dimensional landmark points from the skulls (excluding the distal portion of the beak) from 160 species of birds, suggests that the avian skull has evolved as an integrated unit (Klingenberg and Marugán-Lobón, 2013). Similar findings have been suggested using PLS on smaller taxonomic subsets with support recovered for integrated evolution between two subsets of landmarks delimiting the beak from the rest of the skull in analyses on six corvid species (Kulemeyer et al., 2009), on 147 species of birds of prey (Bright et al., 2016), and on 170 species of parrots and cockatoos (Bright et al., 2019).

Finding support for integrated evolution does not necessarily rule out semi-independent evolution between modules. In other words, it is possible for morphological components to evolve in an integrated manner but with certain regions still evolving semi-independently from one another (Drake and Klingenberg, 2010). More recently developed methods (e.g. Goswami and Finarelli 2016; Adams and Collyer 2019) allow us to quantify the strength of covariation between modules under any given hypothesis while allowing for direct comparisons of the support for the different modularity hypotheses. The ability to compare multiple, sometimes similar hypotheses is particularly useful given that, depending on where landmarks are placed, the delineations of what landmarks should be ascribed to a hypothesized module (particularly for landmarks that are placed along borders) are not necessarily clear. To date, studies of bird clades using these methods have yielded highly variable results. Using a high-resolution geometric morphometric dataset with a likelihood-based approach (Evaluating Modularity with Maximum Likelihood: EMMLi, Goswami and Finarelli, 2016), Felice and Goswami (2018) found that, of the 16 different modularity hypotheses assessed, a seven-module hypothesis was best supported across a broad phylogenetic sample of 352 bird species. However, it has recently been noted that EMMLi appears to have high type I error rates, and a tendency to overfit modular hypotheses compared to a newer method for evaluating modularity hypotheses based on an effect size measure (Adams and Collyer, 2019). Given this disagreement on whether avian skull evolution has been characterized by modular evolution or not, it is imperative to reevaluate modular patterns in a nuanced framework.

We used detailed three-dimensional data and recently developed methods to evaluate support for different hypotheses of modularity in an ecologically and morphologically diverse order of birds: the Charadriiformes. Charadriiformes encompasses ⍰370 species of sandpipers, plovers, gulls, and their relatives. Ecological diversity within the order is high and reflected in their skull morphology, especially in the beak (Barbosa and Moreno, 1999). For example, while common ringed plovers (*Charadrius hiaticula*) have small, stout skulls that are adapted to surface pecking, long-billed curlews (*Numenius americanus*) have extremely long and curved skulls that are used to probe soft sediments for prey (del Hoyo et al., 1996), and Atlantic puffins (*Fractercula arctica*) have very deep beaks and wide skulls that they use to capture fish while diving underwater (Lowther et al., 2020). This morphological diversity, combined with the availability of a comprehensive, well-resolved, time-calibrated phylogeny (Černý and Natale, 2022), makes Charadriiformes a good model for studying the evidence for covariation between avian skull components during their evolution. Using data describing skull morphologies of 262 out of 11370 extant species of Charadriiformes, we assessed support for a set of 11 modularity hypotheses at both the ordinal and subgroup level while simultaneously assessing how varying the density of our 3D landmark data impacted our results. We recovered the strongest support for a two-module modularity hypothesis where the beak evolves semi-independently from, but still in a correlated manner with, the rest of the skull. Additionally, we find evidence that the support for this hypothesis is largely driven by one subgroup, with the other subgroup examined exhibiting unique evolutionary dynamics. Our results highlight key methodological considerations and suggest a framework in which future modularity research may operate.

## 3 Materials and Methods

### 3.1 Morphological Data Collection and Visualization

Morphological data was taken from Natale and Slater (2022). This dataset comprises 255 three-dimensional landmarks (42 landmarks placed at discrete locations, 213 semi-landmarks) taken from 262 species (represented by over 400 individual specimens) of Charadriiformes. These landmarks were placed on all portions of the skull (see fig 1), with relatively dense semi-landmarks placed along the sides and tops of the beak to limit the Pinocchio effect (Walker, 2000; Zelditch et al., 2004), where a distant landmark (e.g. tip of the beak) can cause spurious results in analyses. All landmarks were digitized using Stratovan Checkpoint (Stratovan Corporation, 2018) from three-dimensional surface scans that capture the external bony morphology of skulls housed in museum skeletal collections. For further details on scan settings, digitization, and landmark processing, we refer readers to Natale and Slater (2022). A full list of the specimens used in our analysis is available in the Supplementary File in Table 1. All analyses were run in R (RStudio Team, 2019) unless otherwise noted.

**Figure 1:**
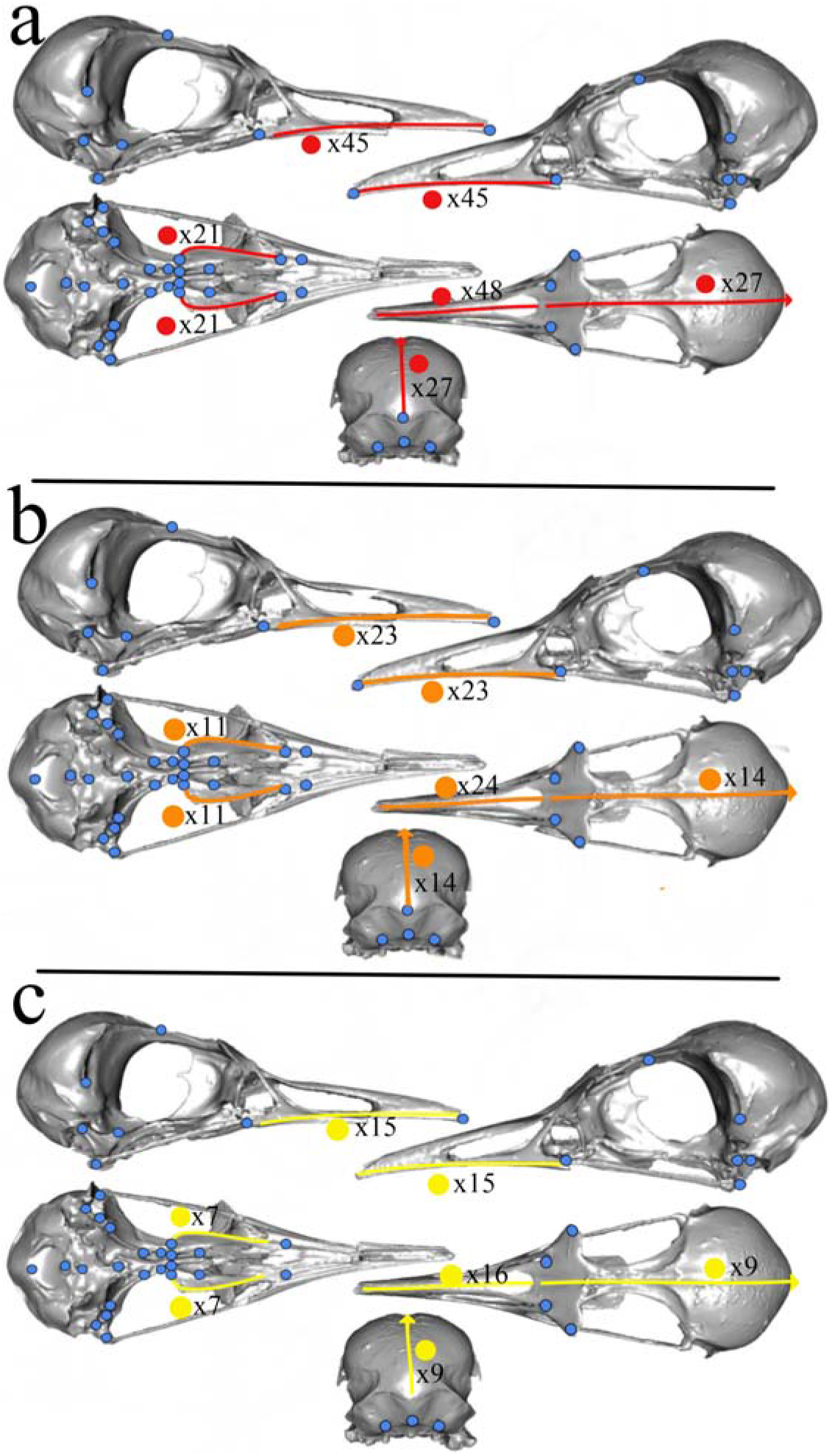
The a) high, b) intermediate, and c) low density landmark schemes used in these analyses depicted on a surface scan of a least tern *Sterna antillarum* (FMNH 376281) skull. Blue dots indicate type I landmarks that are identical under each of the three different densities, with the exception of the low density (panel c) which contains four less type I landmarks. The red, orange, and yellow lines indicate the placement of type II (sliding semi-landmarks) at full (red), intermediate (orange), and low (yellow) densities. The number along each line indicates the number of semi-landmarks placed along that curve under the three distinct densities. A full description of landmarks is available in the Supplementary file, table 2. Figure adapted from Natale and Slater (2022).

**Table 1:** Descriptions of each of the 11 modularity hypotheses compared. The abbreviation used for each hypothesis is given in the lefthand column with the number inside the parenthesis following each name denoting the number of modules in that hypothesis. These descriptions refer to the beak and braincase hypothesis 1 (BKBR1), beak and braincase hypothesis 2 (BKBR2), the FeliceGoswami (FG) hypothesis, and its 8 variations (FG1, FG2, etc.).The righthand column includes hypothesized modules, separated by brackets, with the bones included in each hypothesized module given by distinct letter codes. See figure 3 for the letter codes used here.

| Hypothesis | Modules |
| --- | --- |
| FG (7) | [F,O,Po][Fm][B][Pl][Pt][Q][J,Pm] |
| FG1 (7) | [F(part),O,Po][Fm][B][Pl][Pt][Q][F(part),J,Pm] |
| FG2 (8) | [F(part),O,Po][Fm][B][Pl][Pt][Q][F(part)][J,Pm] |
| FG3 (5) | [F,O,Po,Fm,J,Pm][B][Pl][Pt][Q] |
| FG4 (4) | [F,O,Po,Fm,B,J,Pm][Pl][Pt][Q] |
| FG5 (5) | [F,O,Po,B,J,Pm][Fm][Pl][Pt][Q] |
| FG6 (5) | [F,O,Po][Fm,B,J,Pm][Pl][Pt][Q] |
| FG7 (8) | [F,O][Po][Fm][B][Pl][Pt][Q][J,Pm] |
| FG8 (9) | [F,O][Po][Fm][B][Pl][Pt][Q][J][Pm] |
| BKBR1 (2) | [F,O,Po,Fm,B][Pl,Pt,Q,J,Pm] |
| BKBR2 (2) | [F,O,Po,Fm,B,Pl,Pt,Q,J][Pm] |

**Table 2:** CR scores, P values, and Z*_cr_* scores for each modularity hypothesis tested using the full (FLL), intermediate (INT), and low (LOW) density landmark data. The CR scores indicate the degree of correlation during morphological change among the modules in any given hypothesis. CR values closer to 0 indicate high modularity with sets of landmark points evolving more independently from one another, while CR values closer to 1 indicate low modularity where sets of landmarks tend to evolve in a more covaried manner with one another. P values compare the observed CR to permutations to determine if the evidence for modular evolution under that hypothesis is stronger than expected by chance (p*<*0.05). Z*_cr_* scores represent a measure of support (see fig 5 for statistical comparisons of these values) for that hypothesis with more negative values representing a stronger modular signal under that hypothesis.

| Hypothesis | CR<br>(FLL) | P<br>(FLL) | $Z_{cr}$<br>(FLL) | CR<br>(INT) | P<br>(INT) | $Z_{cr}$<br>(INT) | CR<br>(LOW) | P<br>(LOW) | $Z_{cr}$<br>(LOW) |
| --- | --- | --- | --- | --- | --- | --- | --- | --- | --- |
| FG | 0.82 | 0.001 | -3.50 | 0.83 | 0.001 | -3.88 | 0.83 | 0.001 | -3.34 |
| FG1 | 0.82 | 0.001 | -3.491 | 0.84 | 0.001 | -3.76 | 0.83 | 0.001 | -3.36 |
| FG2 | 0.80 | 0.001 | -4.92 | 0.81 | 0.001 | -4.50 | 0.82 | 0.01 | -3.74 |
| FG3 | 0.78 | 0.001 | -4.92 | 0.79 | 0.001 | -4.64 | 0.80 | 0.01 | -4.19 |
| FG4 | 0.78 | 0.001 | -4.70 | 0.79 | 0.001 | -4.62 | 0.79 | 0.001 | -4.52 |
| FG5 | 0.83 | 0.002 | -3.03 | 0.85 | 0.006 | -2.82 | 0.86 | 0.008 | -2.58 |
| FG6 | 0.77 | 0.001 | -4.80 | 0.78 | 0.001 | -4.54 | 0.78 | 0.001 | -4.36 |
| FG7 | 0.77 | 0.001 | -4.02 | 0.78 | 0.001 | -5.07 | 0.78 | 0.001 | -4.17 |
| FG8 | 0.79 | 0.001 | -4.84 | 0.79 | 0.001 | -5.29 | 0.79 | 0.001 | -4.47 |
| BKBR1 | 0.83 | 0.001 | -14.28 | 0.85 | 0.001 | -8.93 | 0.86 | 0.001 | -5.97 |
| BKBR2 | 0.85 | 0.001 | -19.73 | 0.86 | 0.001 | -11.94 | 0.85 | 0.001 | -7.87 |

Goswami et al. (2019) noted that variation in the density of landmark placement might lead to different outcomes when testing modularity hypotheses. To assess the sensitivity of our results to changes in the density and spread of morphometric data, all ordinal level analyses were conducted on three landmark sets representing three different densities of landmarks (fig 1): The full set of 255 landmarks, an intermediate density set of 154 landmarks (42 landmarks placed at discrete locations, 112 semi-landmarks), and a low density set of 113 landmarks (38 landmarks placed at discrete locations, 75 semi-landmarks). The three distinct densities of landmarks predominantly differed in the number of semi-landmarks placed along the curves (fig 1), but the lowest density landmark set also contained four fewer type I landmarks (see Supplementary File, table 2 for details). Prior to analyses, we slid all semi-landmarks using the slider3d function in the R (RStudio Team, 2019) package Morpho Schlager (2017) and performed Procrustes superimposition using the procSym function, also from the Morpho package, to remove the effects of size, translation, and orientation. To better interpret the results of the following sections and to understand the breadth of charadriiform skull shapes, we also performed a principal components analysis on the Procrustes aligned landmark data for the whole skull using the procSym function in the Morpho package (Schlager, 2017).

### 3.2 Phylogeny

We used the charadriiform phylogeny from Černý and Natale (2022). Full details regarding the inference of this tree are available there. Briefly, the tree was generated from a supermatrix of previously published molecular sequences (12 nuclear, 15 mitochondrial) taken from GenBank (Benson et al., 2013) and a 69-character morphological dataset from Chu (1995). RaXML (Stamatakis, 2014) was used to generate a Maximum Likelihood topology, which was then time-scaled in the program mcmctree (Yang, 2007) using 14 fossil calibrations and a relaxed uncorrelated molecular clock. Prior to our analyses, we removed tips from the tree that were not present in our morphological dataset using the drop.tip function from the R package ape (Paradis et al., 2020).

### 3.3 Modularity Hypotheses

We assessed support for a total of 11 distinct modularity hypotheses (fig 2, see fig 3 for a legend noting the different bones in the avian skull). These hypotheses can be broadly divided into two groups. The first involves variations on the seven-module hypotheses of Felice and Goswami (2018). We did not include the nasal module in this original seven-module hypothesis as its structure cannot be reliably found across the surface scans we used here. Therefore, we instead adopted a similar version to their original Felice and Goswami (2018) model, adjusting for our methodological choices (i.e. differing landmark placements, keeping the quadrate and pterygoid as separate modules). We tested support for this modified hypothesis and 8 variations of it based on the relatively complex, seven-module version by Felice and Goswami (2018). Hereafter, we refer to the first version of this hypothesis as the FeliceGoswami (FG) hypothesis with the variations on this hypothesis labeled sequentially {FG1, FG2, . . ., FG8}. Secondly, we assessed support for two variations of a relatively simpler ‘beak and braincase model’ (hereafter referred to as the BKBR1 and BKBR2 hypotheses) that have been suggested in previous work (e.g., Bright et al. 2016). Variations on both of these hypotheses were generated by altering groupings of landmarks based on either knowledge of the functional interactions (e.g. the quadrate does not touch the palate bones, but is functionally linked during mouth opening and closing; Olsen and Westneat 2016) or proximity between different parts of the skull (e.g. the basioccipital bone is separated from the neighboring parietal bone in some hypotheses). In addition, we compared support for these 11 hypotheses to a null hypothesis, where all landmarks are considered part of one single module (i.e. no modularity).

**Figure 2:**
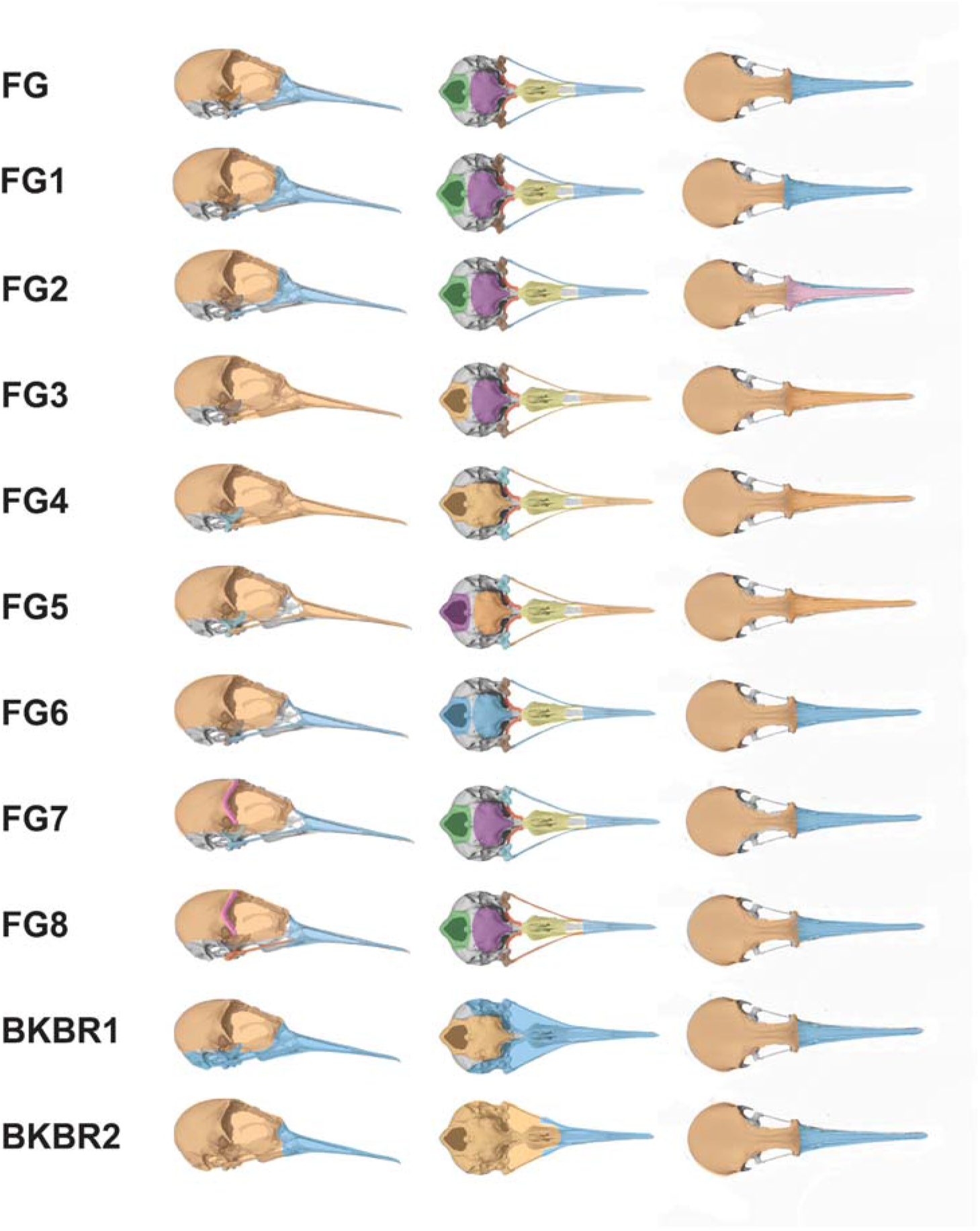
Depictions of the 11 different hypotheses of charadiiform modular skull evolution shown on a skull of a red knot, *Calidris canutus* (FMNH 362893). In each image, distinct colors represent distinct modules. See table 1 for details on each modularity hypothesis. The null model of no modularity is not shown here.

**Figure 3:**
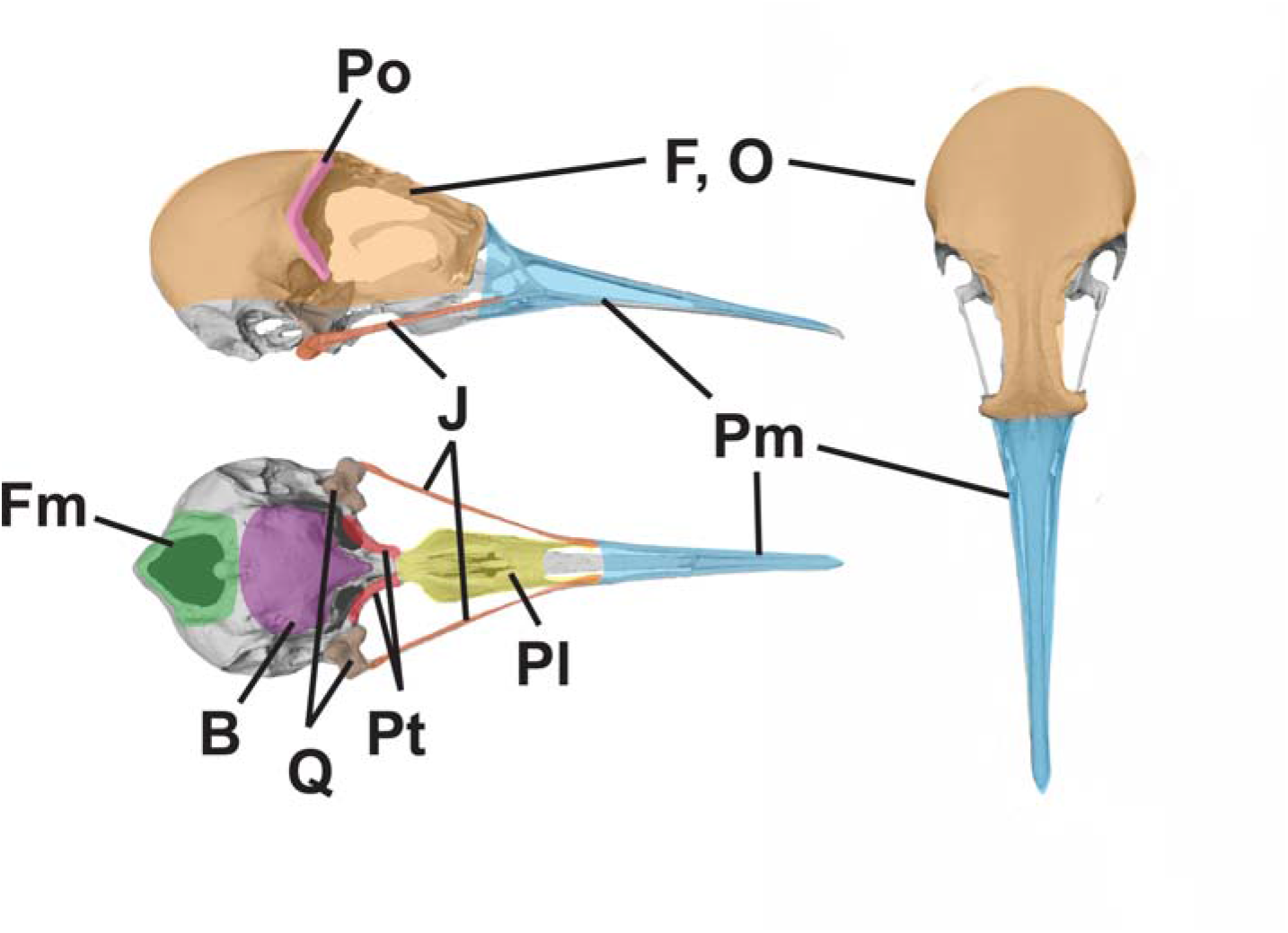
A legend of all the bones used in our analysis shown on a skull of a red knot, *Calidris canutus* (FMNH 362893). The letter codes refer to the following bones: F: frontal, O: occipital, Po: postorbital, Fm: foramen magnum, B: basisphenoid, Pl: palatine, Pt: pterygoid, Q: quadrate, J: jugal, Pm: premaxillary.

### 3.4 Testing Support for Distinct Hypotheses of Modular Evolution Across all Charadriiformes

We used the Covariance Ratio (CR), as computed in the phylo.modularity function from the R package geomorph (Adams et al., 2021; Baken et al., 2021), to quantify the degree of the modular signal under each of the 11 different hypotheses. CR quantifies modular signal by determining and comparing the covariances within and between pre-defined modules (Adams and Collyer, 2019). Typically, CR varies between 0 (no covariation between modules but high covariation within modules; high modularity) and 1 (strong covariation between modules relative to between modules; low modularity), though it may theoretically exceed 1 if covariances between modules are greater than covariances within them (Adams and Collyer, 2019). When there are more than two modules in a given hypothesis, the CR value is averaged across pairwise comparisons of modules. A permutation procedure is then used to determine if the modular signal in the data under each distinct hypothesis is stronger than expected by chance (Adams, 2016). For each analysis, we used 999 permutations to evaluate significance.

The phylo.modularity function also returns the standardized effect sizes Z_CR_ (Adams and Collyer, 2019) for each hypothesis which we used to compare support for the 11 modularity hypotheses. Z_CR_ evaluates the magnitude of the observed effect relative to the random outcomes of the permutation procedure used to evaluate the significance of the CR. Paired effect sizes, 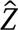_12_, were also computed using the compare.CR function in the geomorph package (Adams et al., 2021) to allow direct comparisons of support between pairs of modularity hypotheses, including the null hypothesis of no modularity (Adams and Collyer, 2019). The compare.CR function assesses support for each hypothesis relative to one another by performing two sample Z tests on the standard error from the results of the CR permutations and will return two sample Z scores and a p-value for each pairwise comparison, indicating whether there is significantly distinct support for any given hypothesis relative to another.

### 3.5 Testing for Variable Patterns of Modular Evolution Across Charadriiform Subgroups

Charadriiformes is a relatively old order of birds (Černý and Natale, 2022) exhibiting considerable ecological diversity (del Hoyo et al., 1996). Recognizing this variation, we hypothesized that limiting tests of modular patterns to the level of the order could mask distinct patterns of modular evolution if present within subgroups. To test this hypothesis, we assessed support using the same CR ratio method described above for the same 11 modularity hypothesis to two subgroups: 1) the Lari (n=114); and, 2) a combined subgroup of the Charadrii and Scolopaci (hereafter referred to as ‘Charadrii+Scolopaci’; n=143). We refer readers to Supplementary File, table 1 for a detailed list of species in each subgroup. The Lari includes species-rich families such as the Laridae (gulls, terns, and skimmers, n =79) and Alcidae (auks and relatives, n = 21), as well as several small families such as the Turnicidae (buttonquails, n= 3) and the monotypic crab plover (family Dromadidae), while the Charadii+Scolopaci subgroup similarly includes two large families (Charadriidae; plovers and lapwings, n = 45 and Scolopacidae; sandpipers and relatives, n = 65), as well as several smaller (e.g. Burhinidae; thick-knees, n = 7) and monotypic families (e.g. the ibisbill; Ibidorhynchidae). The separation of the three charadriiform sublcades (the Lari, the Charadrii, and the Scolopaci) was done in this way given the ecological similarity of the Charadrii and Scolopaci (the clades that contain the most typical shore and wading birds) relative to the Lari (the clade containing predominantly pelagic and diving birds). We chose to examine modularity only at this level as opposed to, for example, the family level, to ensure that the number of specimens exceeded the number of landmarks in each analysis.

For this reason, subgroup-level analyses were also only run using the lowest-density landmark scheme. To aid in the visualization of these morphological differences between these subgroups, we also plotted the three suborders (the Lari, Scolopaci, and Charadrii) on the morphospace generated by the principal components analysis described above.

## 4 Results

### 4.1 Skull Shape Differences Among Charadriiformes

In our principal components analysis (fig. 4) of the full density landmark set, the first principal component (PC1) explained ⍰70% of the variation in shape. Negative PC1 scores correspond to elongated, narrow skulls, while positive PC1 scores correspond to skulls that are shorter and wider. The second principal component (PC2) explained ⍰3% of the variance. Negative PC2 scores correspond to skulls with a large angle between the beak and braincase, a more rounded braincase with large orbits, and more rostrally placed quadrates and foramen magnum. As PC2 scores increase, the skulls become flattened, such that the angle between the beak and braincase decreases, the braincase becomes flatter, and the quadrates and foramen magnum shift caudally. The three subgroups were not equally spread across this morphospace (visualized in figure 4). The Scolopaci and, to a lesser extent, the Charadrii tended to span the whole range of PC1 values, whereas the Lari tended to be restricted to higher PC1 values. This restriction indicates that species within the Lari tended to have shorter, wider skulls, whereas the other two suborders varied in the length and width of the skull. There was less distinction between the suborders along PC2, although the highest values along this axis were seen in the Lari. In general, there was a higher degree of overlap between the Charadrii and the Scolopaci whereas the Lari overlapped less with either suborder.

**Figure 4:**
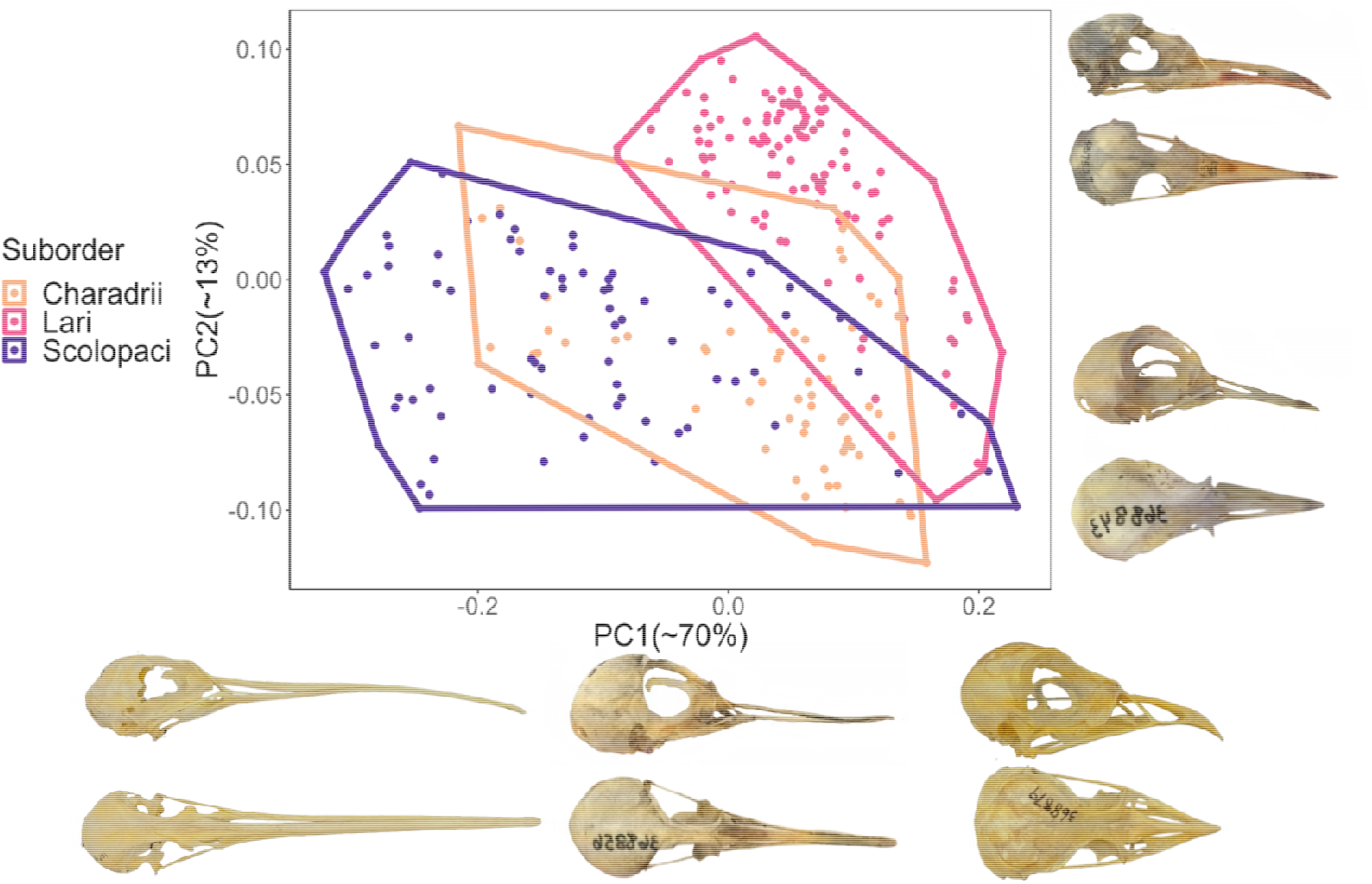
The results of a principal components analysis on 255 3D landmarks from the skulls of 262 different charadriiform species. Each point represents a species morphology digitized from three-dimensional surface scans from 1-2 museum specimens. Each of the three suborders (Charadrii, Lari, and Scolopaci) is encoded by color. Representative examples of changes along each of the first two principal components (explaining about 70 and 13% of overall shape variation, respectively) are shown on the X and Y axes. Species shown along the X axis, listed left to right: long-billed curlew (*Numenius americanus*, FMNH 106289), common sandpiper (*Actitis hypoleucos*, FMNH 368856), and the collared pratincole (*Glareola pratincola*, FMNH 368879). Species along the Y axis, listed top then bottom: Inca tern (*Larosterna inca*, FMNH 437577) and three-banded plover (*Charadrius tricollaris*, FMNH 368844).

### 4.2 Testing Support for Distinct Hypotheses of Modular Evolution Across all Charadriiformes

We found that all eleven hypotheses tested received significant support (p*<*0.05) relative to a null model, although the CR values for all hypotheses and all landmark densities were relatively high (all *>*0.76), indicating that there was a relatively high degree of covariation between modules. The strength of the modular signal under each hypothesis, as indicated by the Z*_cr_* value varied substantially across hypotheses and across differing landmark densities. These Z*_cr_* scores (Adams and Collyer 2019; table 2) indicate that, regardless of the density of the landmark dataset used, the strongest signal of modular evolution was under the BKBR2 hypothesis. This two-module hypothesis separates the beak from the remaining portions of the skull and differs from the BKBR1 hypothesis in that cranial elements that are biomechanically linked to the beak (quadrates and jugals) are retained in the braincase module under the BKBR2 hypothesis. The Z*_cr_* score for the BKBR1 hypothesis always received the second most negative score. Among other hypotheses, the FG5 hypothesis was universally recovered as having the weakest modular signal (as indicated by the least negative Z*_cr_* scores). While the CR values remained relatively static across differing landmark densities, landmark density did relate to the variation in the strength of modular signal over all hypotheses as our densest landmarking scheme returned a higher range of Z*_cr_* scores (16.701) relative to when we used our lowest density landmark set (a range of 4.53).

When support for each of these hypotheses relative to one another was compared via paired effect sizes, we recovered variable results (fig 5). All hypotheses are strongly preferred (p*<*0.05) over the null model of no modularity, regardless of landmark density. When the various versions of the FG hypothesis were compared to one another, very few significant differences were found, the exceptions being support for FG8 over FG4 and FG6 in the intermediate density analyses. Significant support for the two beak and braincase hypotheses, particularly for BKBR2, relative to the FG hypotheses was more frequent but not universal across all landmark densities. Across all densities, distinct support for one beak and braincase hypothesis over the other could not be discriminated (p*>*0.05) on the basis of pairwise effect sizes.

**Figure 5:**
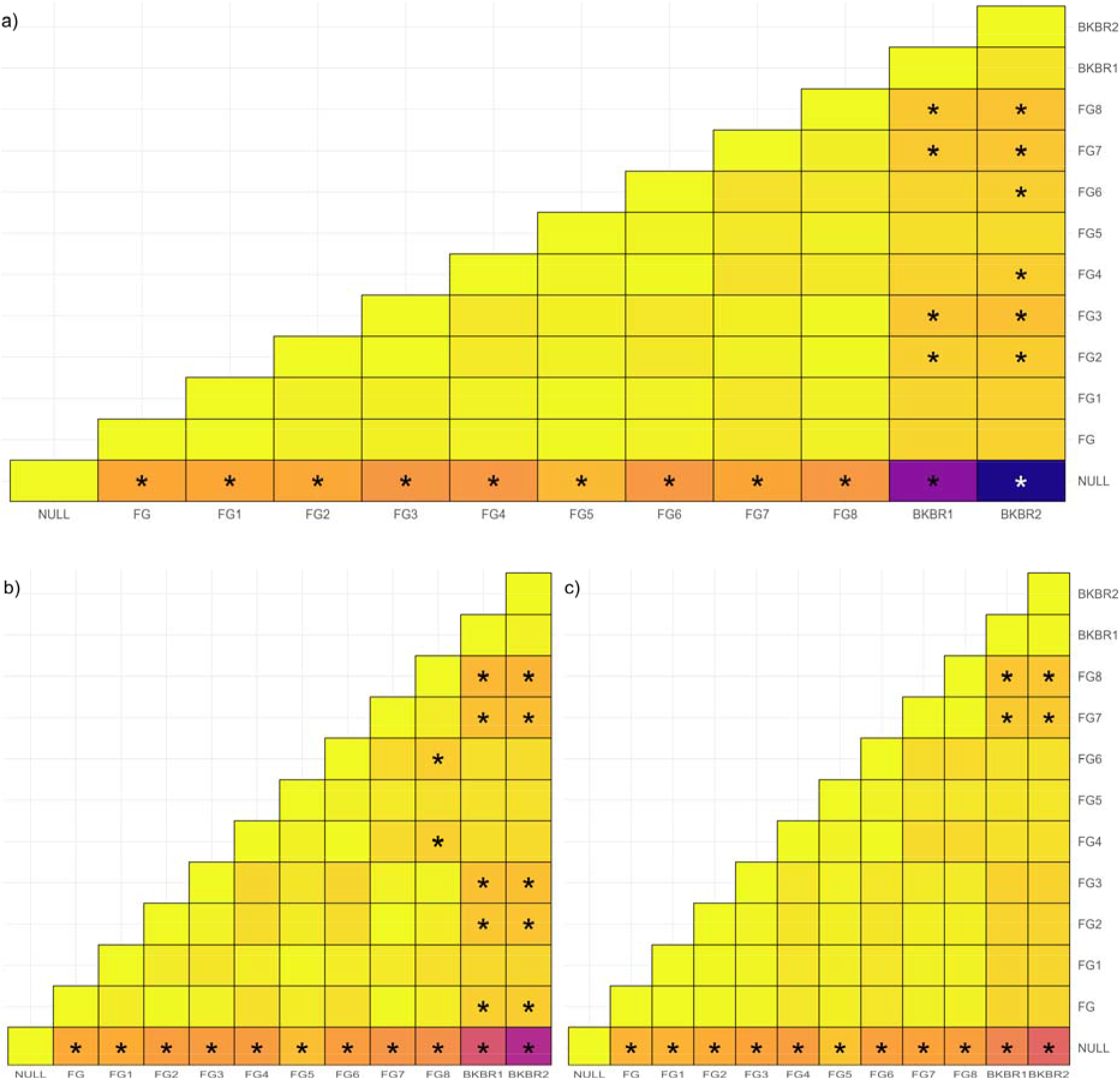
Support at the ordinal level for each modularity hypothesis for charadriiform skull evolution when analyzed using the full (a), intermediate (b), and low (c) density landmark set. Each hypothesis is indicated on the x-axis with an abbreviation (see table 1 and fig 2). The pairwise difference effective size is denoted by the color of each cell with a black or white asterisk (distinct colors are used for increased readability) denoting significantly (p *<* 0.05) different support for that given pair of hypotheses. For these pairwise differences, we determined which hypothesis was favored by comparing the effect sizes (fig 2) with the more negative value indicating the preferred hypothesis. NULL indicates the null model of no modularity to which the others were compared.

### 4.3 Testing for Variable Patterns of Modular Evolution Across Charadriiform Subgroups

We recovered distinct CR values and Z*_cr_* scores for the two subgroups relative to our analyses across all Charadriiformes (fig 6). In the Lari, CR values skewed lower (all *<*0.75) while all CR values skewed higher (all *>* 0.90) in our combined Charadrii+Scolopaci subgroup, suggesting skull modules evolve in a less correlated fashion in the Lari, relative to the Charadrii+Scolopaci. The Z*_cr_*scores for each of the modularity hypotheses also varied between the two subgroups but were again universally more negative (indicating stronger support) in both subgroups relative to a null hypothesis of no modularity. The FG4 hypothesis, in which the upper part of the beak and braincase are treated as a single module, received the least negative value (indicating the weakest support) in both subgroups. The most negative Z*cr* score (indicating the hypothesis with the strongest support) varied between the two families with the BKBR2 hypothesis receiving the most negative scores (similar to our findings at the ordinal level) in the Charadrii+Scolopaci while the FG8 hypothesis, the most finely subdivided of our hypotheses, received the most negative score within the Lari.

**Figure 6:**
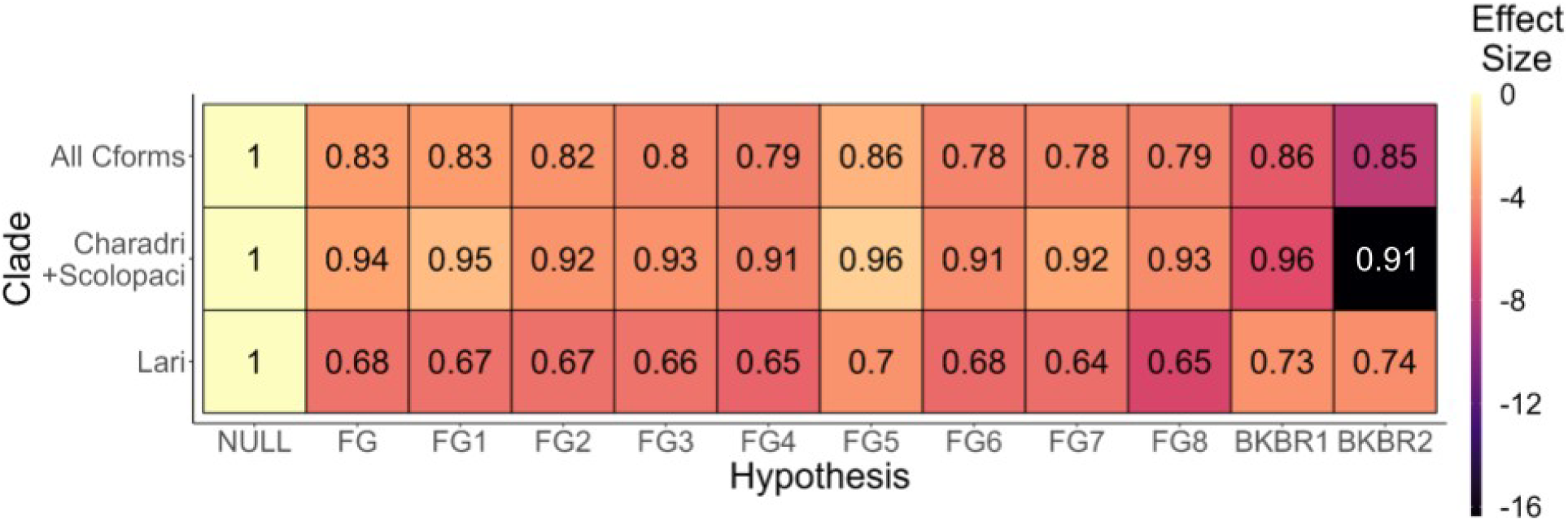
The Covariance Ratio (CR) values and effect sizes (Z*_cr_* scores) for different modularity hypotheses across the charadriiform order (‘All Cforms’) as well within two different charadriiform subgroups: the Lari and a subgroup of the combined suborders Charadrii and Scolopaci (‘Charadrii+ Scolopaci’). The CR value is indicated by the text in each cell, with values closer to 1 indicating a stronger degree of covariation between each pair of modules under that hypothesis. P values for these CR values are not labeled here but we note that all were *<*0.05, indicating significant support for these hypotheses relative to a null model. The effect size (Z*_cr_* scores), given by the color of each cell, demonstrates the relative support for that hypothesis, with more negative values indicating a higher modular signal (and therefore higher support) being detected under that hypothesis. Each hypothesis is indicated on the x-axis with an abbreviation (see table 1 and fig 2). NULL indicates the null model of no modularity to which the others were compared.

Pairwise comparisons of these Z*_cr_* scores revealed complex relationships between support for the hypotheses when compared to one another (fig 7). In both subgroups, all modularity hypotheses were significantly better (p *<* 0.05) than a null model of no modularity, with the exception of the FG5 hypothesis in the Charadii+Scolopaci suborder. Aside from these significant comparisons to the null hypothesis, significant differences in support were (similar to our order level results) recovered most frequently for comparisons between the FG hypotheses and the two versions of the beak and braincase hypothesis. Although the FG8 hypothesis was the most strongly supported within Lari (fig 6, as indicated by the lowest effect size), it is only a significantly more supported hypothesis relative to the FG4 hypothesis for this clade. While the two beak and braincase hypotheses were significantly more supported over a larger number of hypotheses (fig 7A), a significant difference in support for one of the beak and braincase hypotheses relative to the other was not recovered in the Lari (fig 7A). Both the BKBR1 and 2 hypotheses received significantly stronger (p*<*0.05) support than many of the FG hypotheses in the Charadii+Scolopaci (fig 7B), and the two hypotheses were found to be distinctly supported from one another in this subgroup, with the BKBR2 hypothesis receiving a particularly negative Z*_cr_* score of -16.38 (fig 6).

**Figure 7:**
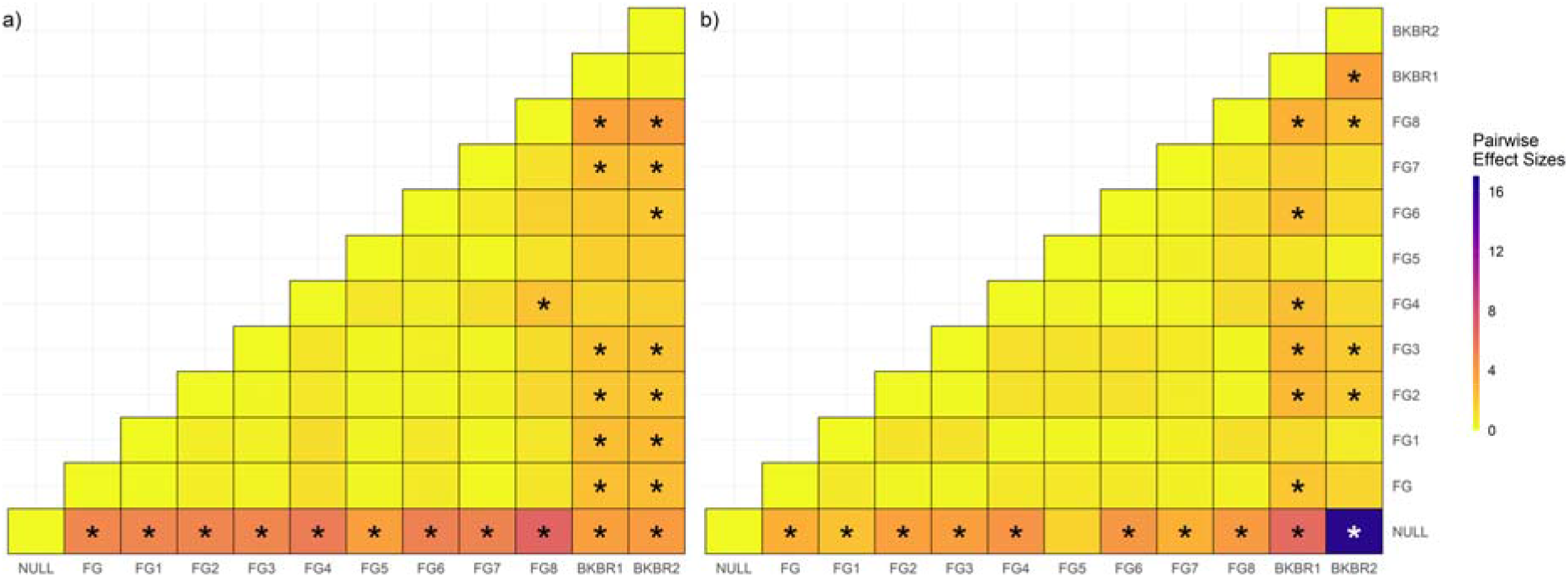
Pairwise effect sizes demonstrating variable support for modularity hypotheses for two different subclades: 1) the Lari and 2) a combined subgroup of the Charadrii + Scolopaci. The pairwise difference is denoted by the color of each cell with a black or white asterisk (distinct colors are used for increased readability) denoting significantly (p *<* 0.05) different support for that given pair of hypotheses. For these pairwise differences, we determined which hypothesis was favored by comparing the effect sizes (table 2) with the more negative value indicating the preferred hypothesis.

## 5 Discussion

In an effort to understand the patterns characterizing the morphological evolution of the vertebrate skull, hypotheses of modular evolution have been tested in a number of clades (Bardua et al., 2019; Adams and Collyer, 2019; Goswami and Finarelli, 2016; Klingenberg and Marugan-Lobon, 2013), using a variety of statistical methods (e.g. Goswami and Finarelli, 2016; Adams and Collyer, 2019). After quantifying the major axes of skull shape variation in a diverse order of birds, the Charadriiformes, we evaluated the support for 11 distinct hypotheses of modular evolution using morphometric data from the skulls of 262 different charadriiform species. We found that, while the density of landmarks used affected the magnitude of the signal detected, the charadriiform skull has likely evolved in a modular manner, with the beak evolving semi-independently from the rest of the skull. Our subgroup-level analyses suggest this broad pattern of evolution is largely driven by one subgroup (the Charadrii+ Scolopaci). Comparisons of this subgroup with another, more morphologically distinct subgroup (the Lari) suggest ecological differences between clades may be driving clade-specific patterns. In general, our results highlight important considerations for analyses of the modular evolution of the avian skull and, by investigating clade-specific results, suggest a framework for future research on the relationship between modularity, morphology, and ecology.

Our strong rejection of a null model of no evolutionary modularity stands in apparent contrast to previous findings based on birds of prey (Bright et al., 2016), parrots (Bright et al., 2019), corvids Kulemeyer et al. (2009), and across a broad sample of all birds (Klingenberg and Marugán-Lobón, 2013) that found evidence for correlated evolution between the beak and the braincase. These studies have relied on methods (Partial Least Squares; Bright et al. 2016, 2019; Kulemeyer et al. 2009 and the RV coefficient; Klingenberg and Marugán-Lobón 2013) that assess the evidence for significant correlations between pre-defined modules. Considering the differing approach of these methods and of that used here (Adams and Collyer, 2019), our results are not incongruous with the idea that the skull has evolved in an inte grated manner as these previous studies have suggested. In agreeance with these previous findings, our analyses all returned relatively high CR values (across all analyses were *>*0.64) indicating a substantial degree of integration between hypothesized modules. However, our results differ from several of these past studies (Bright et al., 2016, 2019; Kulemeyer et al., 2009) in that they include both explicit comparisons of distinct modularity hypotheses relative to one another and a quantifiable measure of the degree of integration between modules, allowing us a more nuanced understanding of the degree of modular evolution occurring. A similar pattern, a degree of modularity with notable integration between modules, was also recovered in a study on domestic dog skull evolution that relied on calculations of RV coefficients, (Drake and Klingenberg, 2010). Drake and Klingenberg (2010) recovered a degree of integration between the face and remaining portions of the skull, but still found significant support for modular evolution between these two elements. It is notable that a methodologically similar study on a broad sample of bird skulls by (Klingenberg and Marugán-Lobón, 2013), found evidence for integration between the beak and braincase as well and rejected a hypothesis of skull modularity. We suggest, given that we found the strongest support for a hypothesis that separated out the beak and braincase, the differing results between our study and that of Klingenberg and Marugán-Lobón (2013) may be explained by their exclusion of the distal portions (roughly corresponding to half) of the beak from their analysis.

Unlike previous studies using methodologies similar to ours that allow for direct comparisons of support for modularity hypotheses, we found little support for complex modularity scenarios. The discrepancy between this finding and that of Felice and Goswami (2018), who suggested that the avian skull is defined by seven evolutionary modules may be explained by the aforementioned tendency of the maximum likelihood method used to favor more complex modularity hypotheses (Adams and Collyer, 2019), a finding that is echoed in research using this method (Goswami and Finarelli, 2016; Mitchell et al., 2021; Marshall et al., 2019; Conith et al., 2020, 2022; Bardua et al., 2020). It is, however, notable that some authors (e.g. Mitchell et al., 2021) have found consistent results between this maximum likelihood method and Covariance Ratio (CR) method, suggesting that in certain species a high number of semi-independent modules may be a genuine pattern. While our finding that reducing the density of semi-landmarks did not typically affect significance testing may suggest landmark density is not of concern, the extreme Z*_cr_* scores found when we used our highest density landmark set suggest extremely dense landmark placement could cause spurious results. This suggestion is in agreement with that of Goswami et al. (2019) who noted that semi-landmarks tend to exaggerate modularity (specifically within region integration and between region modularity). It is possible that the lack of support for higher degrees of modularity in our work, as compared with Felice and Goswami (2018), may reflect the fact that they placed dense semi-landmarks on regions of the skull that we did not. In general, our results highlight the need for future research to carefully consider methodological choices such as landmark placement, density, and subdivision into hypothesized modules as well as the specific hypothesis testing method used.

Our subgroup level results and the spread of the three charadriiform suborders in our morphospace, however, point to levels of taxonomy as another potential reason for variation in past results. The extremely low Z*_cr_* scores for both the BKBR1 (Z*_cr_* score of -8.47) and BKBR2 (Z*_cr_* score of -16.38) hypothesis in our Charadii+Scolopaci subgroup suggest that the species in that group may be the primary drivers of support for these two hypotheses at the ordinal level. Charadii+Scolopaci contain previously studied species where the notable diversity in skull morphology has been tied to foraging ecology (Natale and Slater, 2022). The skulls of both Charadrii and Scolopaci species were found to primarily differ in terms of length and width, often being extremely long and thin (a morphology which is known to facilitate foraging into soft sediments Barbosa and Moreno 1999; Zweers and Gerritsen 1996). It is unsurprising that if beak lengthening is common in the evolution of these species, as suggested by our principal components analysis and by Barbosa and Moreno (1999), our analyses would detect such strong support for the beak evolving as a semi-independent module. However, it is noteworthy that even under both BKBR hypotheses, the CR values were quite high (*>*0.9), suggesting that the modularity between the beak and braincase, while significant, is low. Navalón et al. (2020) note through a modularity analysis on two notable adaptive radiations - the Hawaiian Honeycreepers and the finches of the Galápagos - that integration between modules may also drive adaptive radiation, as has been suggested in theoretical work (Villmoare, 2013). Further analyses explicitly connecting ecology to modularity are needed in the Charadrii+Scolopaci suborder to confirm our hypothesis, but our results here in light of that of Barbosa and Moreno (1999) and Zweers and Gerritsen (1996) suggest that the correlated, but still semi-independent, evolution of the beak with the remaining portions of the skull in this clade may help facilitate diversification of the skull morphology along an axis where length and width change in a correlated manner.

Our results in the Lari subgroup provide an interesting contrast to the Charadii+Scolopaci as we recovered a much less clear signal of support for any given hypothesis within this sub-group. It is notable the CR values recovered were universally lower in the Lari relative to the Charadii+Scolopaci, suggesting that under any given hypothesis, hypothesized modules tended to evolve more independently from one another. The Lari suborder is diverse containing species that differ substantially in ecology and skull morphology, although they tended to occupy more restricted areas of morphospace relative to the Charadrii+Scolopaci. The majority of the suborder comprises species such as gulls, auks, and terns, that tend to forage through a variety of diving behaviors (del Hoyo et al., 1996), but the suborder also contains several small genera such as Turnicidae (buttonquails) and Glareolidae (coursers and pratincoles) that feed terrestrially or aerially. While it is known in this suborder that foraging ecology also relates strongly to skull shape (Natale and Slater, 2022) and there have been species-level analyses of adaptations for various behaviors in both Lari (Zusi, 1962) and in ecologically similar species (e.g. other diving species Eliason et al., 2020), how changes in the morphologies of various skull components such as beak length or eye width may facilitate specific ecological behaviors (e.g. surface diving, pursuit diving, etc…) is not well understood on a macroevolutionary scale. It is possible that a more decoupled evolution of skull components, as suggested by our analysis, may relate to the ecological diversity in this group, but again a more explicit analysis analyzing how modular shape change, ecology, and the breadth of skull morphologies in this order relate to one another is needed. Similar to our analyses and that of Navalón et al. (2020) who performed an in-depth exploration into a hyperdiverse clade, we encourage future covariance-ratio-based research into other avian clades that have been shown to exhibit great variation in foraging ecology and skull morphology such as Order Gruiformes (rails, cranes, and relatives; Livezey 1998)and Procellariiformes (albatrosses, petrels, and relatives; Mazzochi and Carlos 2022). Performing these types of analyses across clades and comparing and highlighting differences between clades, as has been done here, may help detect larger macroevolutionary patterns regarding avian skull shape.

## 6 Conclusion

Contrary to previous research that has suggested the avian evolution has been characterized either by the skull evolving in one integrated unit or as a series of many anatomically distinct modules, we found support for the evolution of the beak and the rest of the skull evolving as two semi-independent modules throughout charadriiform evolution. We found this result to be robust to the use of different density landmark datasets, although our finding that the magnitude of the modular signal detected depended on landmark density highlights this as an important consideration in future research. We found this strong signal of modular evolution appeared to be driven by one subgroup (the Charadii+Scolopaci) which are known to have particularly diverse skull morphologies that relate to differences in foraging behavior. Future analyses investigating these results in light of variation in foraging mode may help elucidate the reasons for variation between subgroups and highlight broad macroevolutionary patterns of avian skull shape evolution. In general, we suggest future modularity research highlights clade-specific dynamics as they relate to differences in the ecology within and between clades.

## Acknowledgements

We would like to thank Alexa Lamprecht, Anna Wisniewski, David Černý, Graham Slater, Jon Nations, Melissa Wood, Mark Webster, Lucas Carneiro, and John Bates for helpful feedback during both analyses and preparation of this manuscript. We additionally thank Graham Slater for access to computers to run analyses as well as for access to the 3D scanners used to collect the initial data used here. This work would not have been possible without previous data collection facilitated financially by the University of Chicago Steiner award and the Climate Change Initiative funded by the Rebecca Susan Buffett Foundation. Additionally, data collection was made possible by the myriad of museum staff that allowed access to collections. We therefore would also like to thank John Bates and Ben Marks at the Field Museum on Natural History, Mark Peck and Santiago Claramunt at the Royal Ontario Museum, Kimball Garrett and Allison Shultz at the Natural History Museum of Los Angeles County, and Christopher Milensky at the Smithsonian National Museum.

## Data Availability Statement

All data used in these analyses as well as the code used to conduct each analysis are available on Dryad (https://doi.org/10.5061/dryad. hmgqnk9n3). These analyses also relied on two previously published datasets available at (https://doi.org/10.5061/dryad.pc866t1p0) and (https://doi.org/10.5061/dryad. xksn02vjc), with the raw surface scan data available on morphosource (morphosource project ID: 00000C909).

