## Supplementary Material for "Correlated evolution of beak and braincase morphology is present only in select bird clades"

### 1 Introduction

This document contains the supplementary information for (insert citation here) which includes two tables of supplementary data. The first contains the information on the specimens used in these analyses and which suborder they are a part of. The second is information on the landmarks collected from each specimen and which were included in the three different density landmark subsets.

### 2 Specimen Information

Table 1: Specimens used in these analyses. The scientific name, which suborder it was assigned to, and the specimen numbers from which data was collected are included. We refer readers to the methods and supplementary information from (Natale and Slater, 2022) for details on the specimens themselves (e.g. the age, sex, location of collection etc... when available) as well as the processing of these specimens to account for missing or damaged skull portions. The surface scans for all specimens from which landmark data was collected are available on the morphosource (morphosource project ID: 00000C909). Under "Specimens Used" FMNH corresponds to the Field Museum of Natural History, ROM corresponds to the Royal Ontario Museum, USNM corresponds to Smithsonian Museum of Natural History, and LACM= Natural History Museum of Los Angeles County.

| Species Name | Suborder | Specimens Used |
|---|---|---|
| Actitis hypoleucos | Scolopaci | FMNH 368856, FMNH 338296 |
| Actitis macularia | Scolopaci | FMNH 466213, FMNH 379170 |
| Actophilornis africanus | Scolopaci | FMNH 368823, FMNH 93434 |
| Aethia cristatella | Lari | LACM 117830, LACM 117834 |
| Aethia psittacula | Lari | LACM 107192, LACM 102621 |
| Aethia pusilla | Lari | USNM 638874, ROM 0150380 |
| Aethia pygmaea | Lari | USNM 344544, LACM 110691 |
| Alca torda | Lari | LACM 103474, FMNH 376712 |
| Alle alle | Lari | FMNH 105265, FMNH 376328 |
| Anarhynchus frontalis | Charadrii | ROM 141502 |
| Anous minutus | Lari | FMNH 346130, FMNH 346127 |
| Anous stolidus | Lari | LACM 117750, FMNH 346102 |
| Aphriza virgata | Scolopaci | USNM 489303 |
| Arenaria interpres | Scolopaci | FMNH 313996 |
| Arenaria melanocephala | Scolopaci | ROM 115678, FMNH 364720 |
| Attagis gayi | Scolopaci | FMNH 105920 |
| Attagis malouinus | Scolopaci | USNM 490853 |
| Bartramia longicauda | Scolopaci | ROM 110807, FMNH 376174 |
| Brachyramphus brevirostris | Lari | USNM 288086, LACM 110518 |
| Brachyramphus marmoratus | Lari | USNM 557617, FMNH 348355 |
| Burhinus bistriatus | Charadrii | FMNH 289831, LACM 87072 |

| Species Name | Suborder | Specimens Used |
|---|---|---|
| Burhinus capensis | Charadrii | LACM 117292, FMNH 390778 |
| Burhinus oedicnemus | Charadrii | FMNH 104449 |
| Burhinus senegalensis | Charadrii | FMNH 368869, USNM 553053 |
| Burhinus superciliaris | Charadrii | LACM 109827, ROM 159572 |
| Burhinus vermiculatus | Charadrii | FMNH 342526, ROM 156978 |
| Calidris acuminata | Scolopaci | USNM 638728, ROM 137943 |
| Calidris alba | Scolopaci | FMNH 483004, FMNH 341863 |
| Calidris alpina | Scolopaci | FMNH 351273, FMNH 376252 |
| Calidris bairdii | Scolopaci | FMNH 105040, FMNH 93225 |
| Calidris canutus | Scolopaci | FMNH 376221, FMNH 363893 |
| Calidris ferruginea | Scolopaci | FMNH 368863 |
| Calidris fuscicollis | Scolopaci | FMNH 317851, FMNH 376240 |
| Calidris himantopus | Scolopaci | LACM 114330, ROM 109502 |
| Calidris maritima | Scolopaci | ROM 0141203, ROM 0150964 |
| Calidris mauri | Scolopaci | FMNH 342546, FMNH 360227 |
| Calidris melanotos | Scolopaci | FMNH 85720, FMNH 105111 |
| Calidris minuta | Scolopaci | FMNH 368861, FMNH 363894 |
| Calidris minutilla | Scolopaci | FMNH 105220, FMNH 105218 |
| Calidris ptilocnemis | Scolopaci | USNM 224073, ROM 122452 |
| Calidris pusilla | Scolopaci | FMNH 290821, FMNH 105215 |
| Calidris ruficollis | Scolopaci | ROM 136257, ROM 0152134 |
| Calidris tenuirostris | Scolopaci | ROM 0152163 |
| Catharacta antarctica | Lari | LACM 112533 |
| Catharacta maccormicki | Lari | USNM 430484, LACM 102398 |
| Catharacta skua | Lari | LACM 102400, ROM 157280 |
| Catoptrophorus semipalmatus | Scolopaci | FMNH 106365, FMNH 106237 |
| Cepphus carbo | Lari | USNM 347755 |
| Cepphus columba | Lari | USNM 612989, LACM 110513 |
| Cepphus grylle | Lari | USNM 623291, ROM 126909 |
| Cerorhinca monocerata | Lari | USNM 347760, ROM 119522 |
| Charadrius alexandrinus | Charadrii | FMNH 317849, FMNH 376156 |
| Charadrius asiaticus | Charadrii | ROM 156945, ROM 122588 |
| Charadrius australis | Charadrii | USNM 553593 |
| Charadrius bicinctus | Charadrii | ROM 122588, ROM 0136248 |
| Charadrius collaris | Charadrii | ROM 158366 |
| Charadrius dubius | Charadrii | ROM 158391 |
| Charadrius falklandicus | Charadrii | ROM 158349, ROM 129581 |
| Charadrius hiaticula | Charadrii | FMNH 368834, FMNH 368835 |
| Charadrius leschenaultii | Charadrii | ROM 156970, USNM 621501 |
| Charadrius marginatus | Charadrii | LACM 90217, FMNH 429392 |
| Charadrius melodus | Charadrii | ROM 15084, ROM 150802 |
| Charadrius modestus | Charadrii | ROM 158361, ROM 158348 |

| Species Name | Suborder | Specimens Used |
|---|---|---|
| Charadrius montanus | Charadrii | LACM 87090, LACM 117334 |
| Charadrius palidus | Charadrii | ROM 156882 |
| Charadrius pecuarius | Charadrii | FMNH 368837, FMNH 368842 |
| Charadrius ruficapillus | Charadrii | ROM 122594, ROM 122621 |
| Charadrius semipalmatus | Charadrii | FMNH 342530, FMNH 106698 |
| Charadrius tricollaris | Charadrii | FMNH 368843, FMNH 368844 |
| Charadrius vociferus | Charadrii | FMNH 390434, FMNH 454807 |
| Charadrius wilsonia | Charadrii | FMNH 360218, FMNH 376171 |
| Chionis albus | Charadrii | USNM 490217, ROM 128919 |
| Chlidonias hybrida | Lari | FMNH 338016 |
| Chlidonias leucopterus | Lari | FMNH 368885, FMNH 368887 |
| Chlidonias nigra | Lari | FMNH 105798 |
| Creagrus furcatus | Lari | USNM 18492, LACM 101442 |
| Cursorius cursor | Lari | USNM 603507, FMNH 368871 |
| Cursorius temminckii | Lari | ROM 156942 |
| Dromas ardeola | Lari | USNM 488404 |
| Elseyornis melanops | Charadrii | ROM 122710, ROM 122711 |
| Esacus magnirostris | Charadrii | USNM 19649, FMNH 104795 |
| Eudromias morinellus | Charadrii | USNM 603466 |
| Fratercula arctica | Lari | ROM 144107, ROM 144101 |
| Fratercula cirrhata | Lari | ROM 119567, FMNH 364740 |
| Fratercula corniculata | Lari | LACM 117861, LACM 117862 |
| Gallinago gallinago | Scolopaci | FMNH 342533, LACM 93280 |
| Gallinago hardwickii | Scolopaci | USNM 612667, ROM 122701 |
| Gallinago media | Scolopaci | USNM 431836 |
| Gallinago megala | Scolopaci | USNM 621521 |
| Gallinago nigripennis | Scolopaci | USNM 48442, ROM 157621 |
| Gallinago paraguaiae | Scolopaci | USNM 622351, ROM 129620 |
| Gallinago stenura | Scolopaci | USNM 557035 |
| Glareola maldivarum | Lari | USNM 19580 |
| Glareola nordmanni | Lari | USNM 430839 |
| Glareola nuchalis | Lari | USNM 347397 |
| Glareola pratincola | Lari | ROM 156964 |
| Gygis alba | Lari | FMNH 314021, FMNH 314022 |
| Haematopus ater | Charadrii | ROM 128856 |
| Haematopus bachmani | Charadrii | ROM 0146602 |
| Haematopus finschi | Charadrii | LACM 117276, ROM 119646 |
| Haematopus fuliginosus | Charadrii | ROM 122613, ROM 122612 |
| Haematopus leucopodus | Charadrii | USNM 490993, ROM 158356 |
| Haematopus moquini | Charadrii | ROM 144605, USNM 558480 |
| Haematopus ostralegus | Charadrii | FMNH 363899, ROM 118045 |
| Haematopus palliatus | Charadrii | USNM 623506, ROM 123861 |
| Haematopus unicolor | Charadrii | ROM 123067, ROM 146618 |

| Species Name | Suborder | Specimens Used |
|---|---|---|
| Heteroscelus brevipes | Scolopaci | ROM 0159634 |
| Heteroscelus incanus | Scolopaci | USNM 612203, LACM 87138 |
| Himantopus himantopus | Charadrii | USNM 431803, ROM 156898 |
| Himantopus melanurus | Charadrii | LACM 109597 |
| Himantopus mexicanus | Charadrii | ROM 114769 |
| Hydrophasianus chirurgus | Scolopaci | USNM 226034 |
| Ibidorhyncha struthersii | Charadrii | USNM 292767 |
| Irediparra gallinacea | Scolopaci | USNM 553360 |
| Jacana jacana | Scolopaci | LACM 87052 |
| Jacana spinosa | Scolopaci | FMNH 105999 |
| Larosterna inca | Lari | FMNH 437527, LACM 107162 |
| Larus argentatus | Lari | FMNH 342553, LACM 117517 |
| Larus atlanticus | Lari | USNM 635816 |
| Larus atricilla | Lari | FMNH 376259, LACM 117558 |
| Larus audouinii | Lari | USNM 488784 |
| Larus bulleri | Lari | ROM 116397, ROM 153422 |
| Larus cachinnans | Lari | LACM 120527, LACM 120528 |
| Larus californicus | Lari | FMNH 105796, ROM 76529 |
| Larus canus | Lari | FMNH 363901, FMNH 398933 |
| Larus cirrocephalus | Lari | LACM 109733, ROM 156972 |
| Larus crassirostris | Lari | USNM 633465 |
| Larus delawarensis | Lari | FMNH 495258, LACM 117501 |
| Larus dominicanus | Lari | USNM 631732, LACM 103871 |
| Larus fuliginosus | Lari | LACM 87185 |
| Larus fuscus | Lari | USNM 631734, USNM 631736 |
| Larus genei | Lari | USNM 500267, LACM 120525 |
| Larus glaucescens | Lari | LACM 100742, USNM 635127 |
| Larus glaucoides | Lari | USNM 637972, ROM 119866 |
| Larus hartlaubii | Lari | USNM 558519, ROM 156844 |
| Larus heermanni | Lari | FMNH 338164, FMNH 338163 |
| Larus hyperboreus | Lari | ROM 0091342, FMNH 105260 |
| Larus ichthyaetus | Lari | USNM 502138 |
| Larus livens | Lari | LACM 103872, LACM 104291 |
| Larus maculipennis | Lari | ROM 129555, ROM 158353 |
| Larus marinus | Lari | FMNH 338219, ROM 91533 |
| Larus melanocephalus | Lari | LACM 120526 |
| Larus minutus | Lari | ROM 124006 |
| Larus modestus | Lari | LACM 110084, ROM 158372 |
| Larus novaehollandiae | Lari | FMNH 390848, ROM 0140609 |
| Larus occidentalis | Lari | FMNH 105459, LACM 101099 |
| Larus pacificus | Lari | ROM 136309, FMNH 338158 |
| Larus philadelphia | Lari | FMNH 376261, ROM 137169 |
| Larus pipixcan | Lari | FMNH 470337, USNM 622198 |

| Species Name | Suborder | Specimens Used |
| --- | --- | --- |
| Larus ridibundus | Lari | LACM 120522, LACM 120523 |
| Larus schistisagus | Lari | USNM 500777 |
| Larus serranus | Lari | USNM 645298, LACM 110060 |
| Larus thayeri | Lari | USNM 347252, ROM 119860 |
| Leucophaeus scoresbii | Lari | USNM 345119, LACM 103815 |
| Limnodromus griseus | Scolopaci | LACM 109401 |
| Limnodromus scolopaceus | Scolopaci | USNM 630671, LACM 117431 |
| Limosa fedoa | Scolopaci | FMNH 23510, USNM 557584 |
| Limosa haemastica | Scolopaci | USNM 489609, ROM 79406 |
| Limosa lapponica | Scolopaci | LACM 87092, USNM 555156 |
| Limosa limosa | Scolopaci | USNM 320129, ROM 157287 |
| Lymnocryptes minimus | Scolopaci | LACM 89975 |
| Metopidius indicus | Scolopaci | USNM 343995 |
| Microparra capensis | Scolopaci | ROM 117587 |
| Numenius americanus | Scolopaci | FMNH 106289, ROM 146877 |
| Numenius arquata | Scolopaci | USNM 553828, FMNH 363888 |
| Numenius madagascariensis | Scolopaci | ROM 127514, USNM 500255 |
| Numenius minutus | Scolopaci | USNM 347648 |
| Numenius phaeopus | Scolopaci | FMNH 385840, USNM 638835 |
| Numenius tahitiensis | Scolopaci | USNM 289233, LACM 103118 |
| Oreopholus ruficollis | Charadrii | FMNH 104122 |
| Pagophila eburnea | Lari | ROM 127323, ROM 147098 |
| Phaetusa simplex | Lari | USNM 345828, FMNH 105560 |
| Phalaropus fulicarius | Scolopaci | LACM 100444, LACM 113112 |
| Phalaropus lobatus | Scolopaci | FMNH 317852, FMNH 106365 |
| Philomachus pugnax | Scolopaci | FMNH 104299, FMNH 104265 |
| Pluvialis apricaria | Scolopaci | ROM 158379 |
| Pluvialis dominica | Charadrii | FMNH 105252, ROM 155330 |
| Pluvialis fulva | Scolopaci | USNM 632112 |
| Pluvialis squatarola | Charadrii | FMNH 376148, LACM 87074 |
| Pluvianus aegyptius | Charadrii | FMNH 379122, FMNH 378793 |
| Ptychoramphus aleuticus | Lari | FMNH 364729, LACM 117815 |
| Recurvirostra americana | Scolopaci | FMNH 364175, FMNH 364177 |
| Recurvirostra andina | Charadrii | ROM 159573 |
| Recurvirostra avosetta | Scolopaci | FMNH 338433, USNM 429088 |
| Rhinoptilus africanus | Lari | USNM 431520, LACM 117298 |
| Rhinoptilus chalcopterus | Lari | USNM 321515, ROM 114357 |
| Rhinoptilus cinctus | Lari | ROM 114395, ROM 114393 |
| Rhodostethia rosea | Lari | USNM 491608, USNM 491609 |
| Rissa brevirostris | Lari | USNM 643045, ROM 150523 |
| Rissa tridactyla | Lari | FMNH 105263, ROM 150453 |
| Rostratula benghalensis | Scolopaci | FMNH 393178, USNM 613014 |

| Species Name | Suborder | Specimens Used |
|---|---|---|
| Rostratula semicollaris | Scolopaci | USNM 612032 |
| Rynchops flavirostris | Lari | FMNH 368889, FMNH 313056 |
| Rynchops niger | Lari | FMNH 398882, ROM 153490 |
| Scolopax minor | Scolopaci | FMNH 438027, LACM 113134 |
| Scolopax rusticola | Scolopaci | USNM 292760, LACM 120516 |
| Steganopus tricolor | Scolopaci | FMNH 106235, FMNH 106234 |
| Stercorarius longicaudus | Lari | FMNH 105258, FMNH 105259 |
| Stercorarius parasiticus | Lari | LACM 87186, LACM 117483 |
| Stercorarius pomarinus | Lari | LACM 117477, FMNH 360238 |
| Sterna albifrons | Lari | FMNH 443603 |
| Sterna aleutica | Lari | USNM 561252 |
| Sterna anaethetus | Lari | FMNH 360250, ROM 118653 |
| Sterna antillarum | Lari | FMNH 376281, LACM 117649 |
| Sterna bengalensis | Lari | USNM 488347 |
| Sterna bergii | Lari | USNM 558526, USNM 559819 |
| Sterna caspia | Lari | FMNH 368884, FMNH 360641 |
| Sterna dougallii | Lari | LACM 92016, FMNH 360247 |
| Sterna elegans | Lari | USNM 612541, LACM 110168 |
| Sterna forsteri | Lari | LACM 117642 |
| Sterna fuscata | Lari | FMNH 364607, FMNH 376277 |
| Sterna hirundinacea | Lari | ROM 129623, LACM 117621 |
| Sterna hirundo | Lari | FMNH 432821, LACM 117627 |
| Sterna lunata | Lari | FMNH 338042, LACM 104037 |
| Sterna maxima | Lari | FMNH 376285, FMNH 379033 |
| Sterna nilotica | Lari | FMNH 368882, FMNH 368883 |
| Sterna paradisaea | Lari | FMNH 317854, LACM 100318 |
| Sterna sandvicensis | Lari | FMNH 313439, LACM 117725 |
| Sterna striata | Lari | USNM 15428, ROM 125713 |
| Sterna sumatrana | Lari | FMNH 104716, FMNH 346071 |
| Sterna superciliaris | Lari | USNM 345825 |
| Sterna trudeaui | Lari | USNM 614631, ROM 159579 |
| Sterna vittata | Lari | ROM 129621 |
| Stiltia isabella | Lari | ROM 149448 |
| Synthliboramphus antiquus | Lari | FMNH 364725, USNM 561925 |
| Synthliboramphus craveri | Lari | LACM 87297, LACM 87296 |
| Synthliboramphus hypoleucus | Lari | USNM 291879, LACM 87299 |
| Thinocorus orbignyianus | Scolopaci | USNM 637907, USNM 645613 |
| Thinocorus rumicivorus | Scolopaci | ROM 128899, ROM 128902 |
| Thinornis rubricollis | Charadrii | ROM 122636, ROM 122635 |
| Tringa flavipes | Scolopaci | FMNH 290820, FMNH 376180 |
| Tringa glareola | Scolopaci | FMNH 368854, FMNH 368852 |
| Tringa melanoleuca | Scolopaci | FMNH 379041 |

| Species Name | Suborder | Specimens Used |
|---|---|---|
| Tringa nebularius | Scolopaci | USNM 620113, ROM 156936 |
| Tringa ochropus | Scolopaci | USNM 345515, FMNH 368850 |
| Tringa solitaria | Scolopaci | FMNH 395650, FMNH 363458 |
| Tringa stagnatilis | Scolopaci | ROM 156883, FMNH 106354 |
| Tringa totanus | Scolopaci | USNM 623276, ROM 157255 |
| Tryngites subruficollis | Scolopaci | ROM 151467, ROM 122461 |
| Turnix suscitator | Lari | FMNH 515266 |
| Turnix sylvaticus | Lari | FMNH 289566, FMNH 291238 |
| Turnix varius | Lari | FMNH 338535 |
| Uria aalge | Lari | FMNH 337986, LACM 100702 |
| Uria lomvia | Lari | USNM 488674, FMNH 105266 |
| Vanellus albiceps | Charadrii | FMNH 313288, FMNH 313055 |
| Vanellus armatus | Charadrii | FMNH 503731, ROM 156886 |
| Vanellus cayanus | Charadrii | FMNH 291736 |
| Vanellus chilensis | Charadrii | LACM 117310, FMNH 106242 |
| Vanellus coronatus | Charadrii | FMNH 368825, LACM 117308 |
| Vanellus crassirostris | Charadrii | USNM 291429 |
| Vanellus duvaucelii | Charadrii | USNM 429213 |
| Vanellus indicus | Charadrii | USNM 645821, USNM 645981 |
| Vanellus leucurus | Charadrii | USNM 645956 |
| Vanellus miles | Charadrii | FMNH 106376, FMNH 105363 |
| Vanellus resplendens | Charadrii | USNM 620756, USNM 645378 |
| Vanellus senegallus | Charadrii | USNM 430408 |
| Vanellus spinosus | Charadrii | FMNH 93433, FMNH 368832 |
| Vanellus tectus | Charadrii | FMNH 104512 |
| Vanellus tricolor | Charadrii | USNM 490657, ROM 111951 |
| Vanellus vanellus | Charadrii | LACM 117304, FMNH 105495 |
| Xema sabini | Lari | USNM 553905 |
| Xenus cinereus | Scolopaci | USNM 633448, ROM 0159633 |

### 3 Landmark Descriptions

The landmark datapoints collected from the species noted above for our analyses. The table contains the number ('Num') used to uniquely identify each landmark, whether or not it was included in the full ('FLL'), intermediate ('INT'), and small ('LOW') density dataset, and the description of that landmark. Inclusion in each of the three different density datasets is noted with an 'x'. Under description, we also included the source with which we took inspiration to include that landmark. If no source is included, there was no direct source from which this landmark point was suggested. This table is adapted with permission from Natale and Slater (2022).

Table 2: Landmarks Used in this Study.

| Num | FLL | INT | LOW | Description |
|---|---|---|---|---|
| 1 | x | x | x | Most anterior point on basisphenoid along the midline (Olsen, 2016) |
| 2 | x | x | x | Most posterior point on basisphenoid along the midline (Olsen, 2016) |
| 3 | x | x | | Medial point along occipital crest (Olsen, 2016) |
| 4 | x | x | x | Most posterior point on foramen magnum along the midline (Olsen, 2016) |
| 5 | x | x | x | Point where jugal meets quadrate, left side (Olsen and Westneat, 2016) |
| 6 | x | x | x | Point where jugal meets quadrate, right side (Olsen and Westneat, 2016) |
| 7 | x | x | x | Point where jugal meets upper beak, right side (Olsen and Westneat, 2016) |
| 8 | x | x | x | Point where jugal meets upper beak, left side (Olsen and Westneat, 2016) |
| 9 | x | x | x | Left quadrate: lateral mandibular condyle (Olsen and Westneat, 2016) |
| 10 | x | x | x | Right quadrate: lateral mandibular condyle (Olsen and Westneat, 2016) |
| 11 | x | x | x | Right quadrate: medial mandibular condyle (Olsen and Westneat, 2016) |
| 12 | x | x | x | Left quadrate: medial mandibular condyle (Olsen and Westneat, 2016) |
| 13 | x | x | x | Left quadrate; lateral condyle of otic process (Olsen and Westneat, 2016) |
| 14 | x | x | x | Right quadrate: lateral condyle of otic process (Olsen and Westneat, 2016) |
| 15 | x | x | x | Right quadrate: medial condyle of otic process (Olsen and Westneat, 2016) |
| 16 | x | x | x | Left quadrate: medial condyle of otic process (Olsen and Westneat, 2016) |
| 17 | x | x | x | Center of occipital condyle at anterior opening of foramen magnum (Olsen, 2016) |
| 18 | x | x | x | Left quadrate: most dorsal part of the orbital process (Olsen and Westneat, 2016) |
| 19 | x | x | x | Right quadrate: most dorsal part of the orbital process (Olsen and Westneat, 2016) |
| 20 | x | x | x | Most posterior point along left opisthotic process (aka paraoccipital process), in groove (Olsen, 2016) |

| Num | FLL | INT | LOW | Description |
|---|---|---|---|---|
| 21 | x | x | x | Most posesterior point along right opisthotic process (aka paraoccipital process), in groove (Olsen, 2016) |
| 22 | x | x | | Left palatine: anterior most point on the lateral crest of the palatine (Olsen and Westneat, 2016) |
| 23 | x | x | | Right palatine: anterior most point on the lateral crest of the palatine (Olsen and Westneat, 2016) |
| 24 | x | x | x | Right palatine: posterior most point on the lateral crest of the palatine (Olsen and Westneat, 2016) |
| 25 | x | x | x | Left palatine: posterior most point on the lateral crest of the palatine (Olsen and Westneat, 2016) |
| 26 | x | x | x | Left palatine; anterior most point on the medial palatine crest (Olsen and Westneat, 2016) |
| 27 | x | x | x | Right palatine; anterior most point on the medial palatine crest (Olsen and Westneat, 2016) |
| 28 | x | x | x | Right palatine; posterior most point on the medial palatine crest (Olsen and Westneat, 2016) |
| 29 | x | x | x | Left palatine; posterior most point on the medial palatine crest (Olsen and Westneat, 2016) |
| 30 | x | x | x | Left side: where palatine meets pterygoid (Olsen and Westneat, 2016) |
| 31 | x | x | x | Right side; where palatine meets pterygoid (Olsen and Westneat, 2016) |
| 32 | x | x | x | Most dorsal point on left orbit |
| 33 | x | x | x | Most dorsal point on left orbit |
| 34 | x | x | x | Most posterior point on right orbit |
| 35 | x | x | x | Most posterior point on left orbit |
| 36 | x | x | x | Point where palatine meets pterygoid, left side (Olsen and Westneat, 2016) |
| 37 | x | x | x | Point where palatine meets pterygoid, right side (Olsen and Westneat, 2016) |
| 38 | x | x | | Most distal point on the beak (Olsen and Westneat, 2016) |
| 39 | x | x | x | Left side, point where beak joins braincase on frontal bones |
| 40 | x | x | x | Right side, point where beak joins braincase on frontal bones |
| 41 | x | x | x | Right side, most rostral point along frontal or prefrontal bones in front of orbit |

| Num | FLL | INT | LOW | Description |
|---|---|---|---|---|
| 42 | x | x | x | Left side, most rostral point along frontal or prefrontal bones in front of orbit |
| 43 | x | x | x | Semilandmarks running along the sagittal plane from the frontal-nasal hinge to the medial point on the occipital crest (Olsen, 2016) |
| 44 | x | | | Semilandmarks running along the sagittal plane from the frontal-nasal hinge to the medial point on the occipital crest (Olsen, 2016) |
| 45 | x | x | | Semilandmarks running along the sagittal plane from the frontal-nasal hinge to the medial point on the occipital crest (Olsen, 2016) |
| 46 | x | | x | Semilandmarks running along the sagittal plane from the frontal-nasal hinge to the medial point on the occipital crest (Olsen, 2016) |
| 47 | x | x | | Semilandmarks running along the sagittal plane from the frontal-nasal hinge to the medial point on the occipital crest (Olsen, 2016) |
| 48 | x | | | Semilandmarks running along the sagittal plane from the frontal-nasal hinge to the medial point on the occipital crest (Olsen, 2016) |
| 49 | x | x | x | Semilandmarks running along the sagittal plane from the frontal-nasal hinge to the medial point on the occipital crest (Olsen, 2016) |
| 50 | x | | | Semilandmarks running along the sagittal plane from the frontal-nasal hinge to the medial point on the occipital crest (Olsen, 2016) |
| 51 | x | x | | Semilandmarks running along the sagittal plane from the frontal-nasal hinge to the medial point on the occipital crest (Olsen, 2016) |
| 52 | x | | x | Semilandmarks running along the sagittal plane from the frontal-nasal hinge to the medial point on the occipital crest (Olsen, 2016) |
| 53 | x | x | | Semilandmarks running along the sagittal plane from the frontal-nasal hinge to the medial point on the occipital crest (Olsen, 2016) |
| 54 | x | | | Semilandmarks running along the sagittal plane from the frontal-nasal hinge to the medial point on the occipital crest (Olsen, 2016) |
| 55 | x | x | x | Semilandmarks running along the sagittal plane from the frontal-nasal hinge to the medial point on the occipital crest (Olsen, 2016) |
| 56 | x | | | Semilandmarks running along the sagittal plane from the frontal-nasal hinge to the medial point on the occipital crest (Olsen, 2016) |

| Num | FLL | INT | LOW | Description |
|---|---|---|---|---|
| 57 | x | x | | Semilandmarks running along the sagittal plane from the frontal-nasal hinge to the medial point on the occipital crest (Olsen, 2016) |
| 58 | x | | x | Semilandmarks running along the sagittal plane from the frontal-nasal hinge to the medial point on the occipital crest (Olsen, 2016) |
| 59 | x | x | | Semilandmarks running along the sagittal plane from the frontal-nasal hinge to the medial point on the occipital crest (Olsen, 2016) |
| 60 | x | | | Semilandmarks running along the sagittal plane from the frontal-nasal hinge to the medial point on the occipital crest (Olsen, 2016) |
| 61 | x | x | x | Semilandmarks running along the sagittal plane from the frontal-nasal hinge to the medial point on the occipital crest (Olsen, 2016) |
| 62 | x | | | Semilandmarks running along the sagittal plane from the frontal-nasal hinge to the medial point on the occipital crest (Olsen, 2016) |
| 63 | x | x | | Semilandmarks running along the sagittal plane from the frontal-nasal hinge to the medial point on the occipital crest (Olsen, 2016) |
| 64 | x | | x | Semilandmarks running along the sagittal plane from the frontal-nasal hinge to the medial point on the occipital crest (Olsen, 2016) |
| 65 | x | x | | Semilandmarks running along the sagittal plane from the frontal-nasal hinge to the medial point on the occipital crest (Olsen, 2016) |
| 66 | x | | | Semilandmarks running along the sagittal plane from the frontal-nasal hinge to the medial point on the occipital crest (Olsen, 2016) |
| 67 | x | x | x | Semilandmarks running along the sagittal plane from the frontal-nasal hinge to the medial point on the occipital crest (Olsen, 2016) |
| 68 | x | | | Semilandmarks running along the sagittal plane from the frontal-nasal hinge to the medial point on the occipital crest (Olsen, 2016) |
| 69 | x | x | | Semilandmarks running along the sagittal plane from the frontal-nasal hinge to the medial point on the occipital crest (Olsen, 2016) |
| 70 | x | x | x | Semilandmarks running along the sagittal plane from the frontal-nasal hinge to the medial point on the occipital crest (Olsen, 2016) |
| 71 | x | x | x | Semilandmarks running along proximal curve on the left palatine (Olsen, 2016) |

| Num | FLL | INT | LOW | Description |
|---|---|---|---|---|
| 72 | x | | | Semilandmarks running along proximal curve on the left palatine (Olsen, 2016) |
| 73 | x | x | | Semilandmarks running along proximal curve on the left palatine (Olsen, 2016) |
| 74 | x | | x | Semilandmarks running along proximal curve on the left palatine (Olsen, 2016) |
| 75 | x | x | | Semilandmarks running along proximal curve on the left palatine (Olsen, 2016) |
| 76 | x | | | Semilandmarks running along proximal curve on the left palatine (Olsen, 2016) |
| 77 | x | x | x | Semilandmarks running along proximal curve on the left palatine (Olsen, 2016) |
| 78 | x | | | Semilandmarks running along proximal curve on the left palatine (Olsen, 2016) |
| 79 | x | x | | Semilandmarks running along proximal curve on the left palatine (Olsen, 2016) |
| 80 | x | | x | Semilandmarks running along proximal curve on the left palatine (Olsen, 2016) |
| 81 | x | x | | Semilandmarks running along proximal curve on the left palatine (Olsen, 2016) |
| 82 | x | | | Semilandmarks running along proximal curve on the left palatine (Olsen, 2016) |
| 83 | x | x | x | Semilandmarks running along proximal curve on the left palatine (Olsen, 2016) |
| 84 | x | | | Semilandmarks running along proximal curve on the left palatine (Olsen, 2016) |
| 85 | x | x | | Semilandmarks running along proximal curve on the left palatine (Olsen, 2016) |
| 86 | x | | x | Semilandmarks running along proximal curve on the left palatine (Olsen, 2016) |
| 87 | x | x | | Semilandmarks running along proximal curve on the left palatine (Olsen, 2016) |
| 88 | x | | | Semilandmarks running along proximal curve on the left palatine (Olsen, 2016) |
| 89 | x | x | x | Semilandmarks running along proximal curve on the left palatine (Olsen, 2016) |
| 90 | x | | | Semilandmarks running along proximal curve on the left palatine (Olsen, 2016) |
| 91 | x | x | | Semilandmarks running along proximal curve on the left palatine (Olsen, 2016) |
| 92 | x | x | x | Semilandmarks running along proximal curve on the left palatine (Olsen, 2016) |
| 93 | x | x | x | Semilandmarks running along proximal curve on the right palatine (Olsen, 2016) |

| Num | FLL | INT | LOW | Description |
|---|---|---|---|---|
| 94 | x | | | Semilandmarks running along proximal curve on the right palatine (Olsen, 2016) |
| 95 | x | x | | Semilandmarks running along proximal curve on the right palatine (Olsen, 2016) |
| 96 | x | | x | Semilandmarks running along proximal curve on the right palatine (Olsen, 2016) |
| 97 | x | x | | Semilandmarks running along proximal curve on the right palatine (Olsen, 2016) |
| 98 | x | | | Semilandmarks running along proximal curve on the right palatine (Olsen, 2016) |
| 99 | x | x | x | Semilandmarks running along proximal curve on the right palatine (Olsen, 2016) |
| 100 | x | | | Semilandmarks running along proximal curve on the right palatine (Olsen, 2016) |
| 101 | x | x | | Semilandmarks running along proximal curve on the right palatine (Olsen, 2016) |
| 102 | x | | x | Semilandmarks running along proximal curve on the right palatine (Olsen, 2016) |
| 103 | x | x | | Semilandmarks running along proximal curve on the right palatine (Olsen, 2016) |
| 104 | x | | | Semilandmarks running along proximal curve on the right palatine (Olsen, 2016) |
| 105 | x | x | x | Semilandmarks running along proximal curve on the right palatine (Olsen, 2016) |
| 106 | x | | | Semilandmarks running along proximal curve on the right palatine (Olsen, 2016) |
| 107 | x | x | | Semilandmarks running along proximal curve on the right palatine (Olsen, 2016) |
| 108 | x | | x | Semilandmarks running along proximal curve on the right palatine (Olsen, 2016) |
| 109 | x | x | | Semilandmarks running along proximal curve on the right palatine (Olsen, 2016) |
| 110 | x | | | Semilandmarks running along proximal curve on the right palatine (Olsen, 2016) |
| 111 | x | x | x | Semilandmarks running along proximal curve on the right palatine (Olsen, 2016) |
| 112 | x | | | Semilandmarks running along proximal curve on the right palatine (Olsen, 2016) |
| 113 | x | x | | Semilandmarks running along proximal curve on the right palatine (Olsen, 2016) |
| 114 | x | x | x | Semilandmarks running along proximal curve on the right palatine (Olsen, 2016) |
| 115 | x | x | x | Semilandmarks running along top of the beak (Olsen, 2016) |

| Num | FLL | INT | LOW | Description |
|---|---|---|---|---|
| 116 | x | | | Semilandmarks running along top of the beak (Olsen, 2016) |
| 117 | x | x | | Semilandmarks running along top of the beak (Olsen, 2016) |
| 118 | x | | x | Semilandmarks running along top of the beak (Olsen, 2016) |
| 119 | x | x | | Semilandmarks running along top of the beak (Olsen, 2016) |
| 120 | x | | | Semilandmarks running along top of the beak (Olsen, 2016) |
| 121 | x | x | x | Semilandmarks running along top of the beak (Olsen, 2016) |
| 122 | x | | | Semilandmarks running along top of the beak (Olsen, 2016) |
| 123 | x | x | | Semilandmarks running along top of the beak (Olsen, 2016) |
| 124 | x | | x | Semilandmarks running along top of the beak (Olsen, 2016) |
| 125 | x | x | | Semilandmarks running along top of the beak (Olsen, 2016) |
| 126 | x | | | Semilandmarks running along top of the beak (Olsen, 2016) |
| 127 | x | x | x | Semilandmarks running along top of the beak (Olsen, 2016) |
| 128 | x | | | Semilandmarks running along top of the beak (Olsen, 2016) |
| 129 | x | x | | Semilandmarks running along top of the beak (Olsen, 2016) |
| 130 | x | | x | Semilandmarks running along top of the beak (Olsen, 2016) |
| 131 | x | x | | Semilandmarks running along top of the beak (Olsen, 2016) |
| 132 | x | | | Semilandmarks running along top of the beak (Olsen, 2016) |
| 133 | x | x | x | Semilandmarks running along top of the beak (Olsen, 2016) |
| 134 | x | | | Semilandmarks running along top of the beak (Olsen, 2016) |
| 135 | x | x | | Semilandmarks running along top of the beak (Olsen, 2016) |
| 136 | x | | x | Semilandmarks running along top of the beak (Olsen, 2016) |
| 137 | x | x | | Semilandmarks running along top of the beak (Olsen, 2016) |

| Num | FLL | INT | LOW | Description |
|---|---|---|---|---|
| 138 | x | | | Semilandmarks running along top of the beak (Olsen, 2016) |
| 139 | x | x | x | Semilandmarks running along top of the beak (Olsen, 2016) |
| 140 | x | | | Semilandmarks running along top of the beak (Olsen, 2016) |
| 141 | x | x | | Semilandmarks running along top of the beak (Olsen, 2016) |
| 142 | x | | x | Semilandmarks running along top of the beak (Olsen, 2016) |
| 143 | x | x | | Semilandmarks running along top of the beak (Olsen, 2016) |
| 144 | x | | | Semilandmarks running along top of the beak (Olsen, 2016) |
| 145 | x | x | x | Semilandmarks running along top of the beak (Olsen, 2016) |
| 146 | x | | | Semilandmarks running along top of the beak (Olsen, 2016) |
| 147 | x | x | | Semilandmarks running along top of the beak (Olsen, 2016) |
| 148 | x | | x | Semilandmarks running along top of the beak (Olsen, 2016) |
| 149 | x | x | | Semilandmarks running along top of the beak (Olsen, 2016) |
| 150 | x | | | Semilandmarks running along top of the beak (Olsen, 2016) |
| 151 | x | x | x | Semilandmarks running along top of the beak (Olsen, 2016) |
| 152 | x | | | Semilandmarks running along top of the beak (Olsen, 2016) |
| 153 | x | x | | Semilandmarks running along top of the beak (Olsen, 2016) |
| 154 | x | | x | Semilandmarks running along top of the beak (Olsen, 2016) |
| 155 | x | x | | Semilandmarks running along top of the beak (Olsen, 2016) |
| 156 | x | | | Semilandmarks running along top of the beak (Olsen, 2016) |
| 157 | x | x | x | Semilandmarks running along top of the beak (Olsen, 2016) |
| 158 | x | | | Semilandmarks running along top of the beak (Olsen, 2016) |
| 159 | x | x | | Semilandmarks running along top of the beak (Olsen, 2016) |

| Num | FLL | INT | LOW | Description |
|---|---|---|---|---|
| 160 | x | | x | Semilandmarks running along top of the beak (Olsen, 2016) |
| 161 | x | x | | Semilandmarks running along top of the beak (Olsen, 2016) |
| 162 | x | | | Semilandmarks running along top of the beak (Olsen, 2016) |
| 163 | x | x | x | Semilandmarks running along top of the beak (Olsen, 2016) |
| 164 | x | x | x | Semilandmarks running along the right side of the beak (Olsen, 2016) |
| 165 | x | | | Semilandmarks running along the right side of the beak (Olsen, 2016) |
| 166 | x | x | | Semilandmarks running along the right side of the beak (Olsen, 2016) |
| 167 | x | | x | Semilandmarks running along the right side of the beak (Olsen, 2016) |
| 168 | x | x | | Semilandmarks running along the right side of the beak (Olsen, 2016) |
| 169 | x | | | Semilandmarks running along the right side of the beak (Olsen, 2016) |
| 170 | x | x | x | Semilandmarks running along the right side of the beak (Olsen, 2016) |
| 171 | x | | | Semilandmarks running along the right side of the beak (Olsen, 2016) |
| 172 | x | x | | Semilandmarks running along the right side of the beak (Olsen, 2016) |
| 173 | x | | x | Semilandmarks running along the right side of the beak (Olsen, 2016) |
| 174 | x | x | | Semilandmarks running along the right side of the beak (Olsen, 2016) |
| 175 | x | | | Semilandmarks running along the right side of the beak (Olsen, 2016) |
| 176 | x | x | x | Semilandmarks running along the right side of the beak (Olsen, 2016) |
| 177 | x | | | Semilandmarks running along the right side of the beak (Olsen, 2016) |
| 178 | x | x | | Semilandmarks running along the right side of the beak (Olsen, 2016) |
| 179 | x | | x | Semilandmarks running along the right side of the beak (Olsen, 2016) |
| 180 | x | x | | Semilandmarks running along the right side of the beak (Olsen, 2016) |
| 181 | x | | | Semilandmarks running along the right side of the beak (Olsen, 2016) |

| Num | FLL | INT | LOW | Description |
|---|---|---|---|---|
| 182 | x | x | x | Semilandmarks running along the right side of the beak (Olsen, 2016) |
| 183 | x | | | Semilandmarks running along the right side of the beak (Olsen, 2016) |
| 184 | x | x | | Semilandmarks running along the right side of the beak (Olsen, 2016) |
| 185 | x | | x | Semilandmarks running along the right side of the beak (Olsen, 2016) |
| 186 | x | x | | Semilandmarks running along the right side of the beak (Olsen, 2016) |
| 187 | x | | | Semilandmarks running along the right side of the beak (Olsen, 2016) |
| 188 | x | x | x | Semilandmarks running along the right side of the beak (Olsen, 2016) |
| 189 | x | | | Semilandmarks running along the right side of the beak (Olsen, 2016) |
| 190 | x | x | | Semilandmarks running along the right side of the beak (Olsen, 2016) |
| 191 | x | | x | Semilandmarks running along the right side of the beak (Olsen, 2016) |
| 192 | x | x | | Semilandmarks running along the right side of the beak (Olsen, 2016) |
| 193 | x | | | Semilandmarks running along the right side of the beak (Olsen, 2016) |
| 194 | x | x | x | Semilandmarks running along the right side of the beak (Olsen, 2016) |
| 195 | x | | | Semilandmarks running along the right side of the beak (Olsen, 2016) |
| 196 | x | x | | Semilandmarks running along the right side of the beak (Olsen, 2016) |
| 197 | x | | x | Semilandmarks running along the right side of the beak (Olsen, 2016) |
| 198 | x | x | | Semilandmarks running along the right side of the beak (Olsen, 2016) |
| 199 | x | | | Semilandmarks running along the right side of the beak (Olsen, 2016) |
| 200 | x | x | x | Semilandmarks running along the right side of the beak (Olsen, 2016) |
| 201 | x | | | Semilandmarks running along the right side of the beak (Olsen, 2016) |
| 202 | x | x | | Semilandmarks running along the right side of the beak (Olsen, 2016) |
| 203 | x | | x | Semilandmarks running along the right side of the beak (Olsen, 2016) |

| Num | FLL | INT | LOW | Description |
|---|---|---|---|---|
| 204 | x | x | | Semilandmarks running along the right side of the beak (Olsen, 2016) |
| 205 | x | | | Semilandmarks running along the right side of the beak (Olsen, 2016) |
| 206 | x | x | x | Semilandmarks running along the right side of the beak (Olsen, 2016) |
| 207 | x | | | Semilandmarks running along the right side of the beak (Olsen, 2016) |
| 208 | x | x | | Semilandmarks running along the right side of the beak (Olsen, 2016) |
| 209 | x | x | x | Semilandmarks running along the right side of the beak (Olsen, 2016) |
| 210 | x | x | x | Semilandmarks running along the left side of the beak (Olsen, 2016) |
| 211 | x | | | Semilandmarks running along the left side of the beak (Olsen, 2016) |
| 212 | x | x | | Semilandmarks running along the left side of the beak (Olsen, 2016) |
| 213 | x | | x | Semilandmarks running along the left side of the beak (Olsen, 2016) |
| 214 | x | x | | Semilandmarks running along the left side of the beak (Olsen, 2016) |
| 215 | x | | | Semilandmarks running along the left side of the beak (Olsen, 2016) |
| 216 | x | x | x | Semilandmarks running along the left side of the beak (Olsen, 2016) |
| 217 | x | | | Semilandmarks running along the left side of the beak (Olsen, 2016) |
| 218 | x | x | | Semilandmarks running along the left side of the beak (Olsen, 2016) |
| 219 | x | | x | Semilandmarks running along the left side of the beak (Olsen, 2016) |
| 220 | x | x | | Semilandmarks running along the left side of the beak (Olsen, 2016) |
| 221 | x | | | Semilandmarks running along the left side of the beak (Olsen, 2016) |
| 222 | x | x | x | Semilandmarks running along the left side of the beak (Olsen, 2016) |
| 223 | x | | | Semilandmarks running along the left side of the beak (Olsen, 2016) |
| 224 | x | x | | Semilandmarks running along the left side of the beak (Olsen, 2016) |
| 225 | x | | x | Semilandmarks running along the left side of the beak (Olsen, 2016) |

| Num | FLL | INT | LOW | Description |
|---|---|---|---|---|
| 226 | x | x | | Semilandmarks running along the left side of the beak (Olsen, 2016) |
| 227 | x | | | Semilandmarks running along the left side of the beak (Olsen, 2016) |
| 228 | x | x | x | Semilandmarks running along the left side of the beak (Olsen, 2016) |
| 229 | x | | | Semilandmarks running along the left side of the beak (Olsen, 2016) |
| 230 | x | x | | Semilandmarks running along the left side of the beak (Olsen, 2016) |
| 231 | x | | x | Semilandmarks running along the left side of the beak (Olsen, 2016) |
| 232 | x | x | | Semilandmarks running along the left side of the beak (Olsen, 2016) |
| 233 | x | | | Semilandmarks running along the left side of the beak (Olsen, 2016) |
| 234 | x | x | x | Semilandmarks running along the left side of the beak (Olsen, 2016) |
| 235 | x | | | Semilandmarks running along the left side of the beak (Olsen, 2016) |
| 236 | x | x | | Semilandmarks running along the left side of the beak (Olsen, 2016) |
| 237 | x | | x | Semilandmarks running along the left side of the beak (Olsen, 2016) |
| 238 | x | x | | Semilandmarks running along the left side of the beak (Olsen, 2016) |
| 239 | x | | | Semilandmarks running along the left side of the beak (Olsen, 2016) |
| 240 | x | x | x | Semilandmarks running along the left side of the beak (Olsen, 2016) |
| 241 | x | | | Semilandmarks running along the left side of the beak (Olsen, 2016) |
| 242 | x | x | | Semilandmarks running along the left side of the beak (Olsen, 2016) |
| 243 | x | | x | Semilandmarks running along the left side of the beak (Olsen, 2016) |
| 244 | x | x | | Semilandmarks running along the left side of the beak (Olsen, 2016) |
| 245 | x | | | Semilandmarks running along the left side of the beak (Olsen, 2016) |
| 246 | x | x | x | Semilandmarks running along the left side of the beak (Olsen, 2016) |
| 247 | x | | | Semilandmarks running along the left side of the beak (Olsen, 2016) |

| Num | FLL | INT | LOW | Description |
|---|---|---|---|---|
| 248 | x | x | | Semilandmarks running along the left side of the beak (Olsen, 2016) |
| 249 | x | | x | Semilandmarks running along the left side of the beak (Olsen, 2016) |
| 250 | x | x | | Semilandmarks running along the left side of the beak (Olsen, 2016) |
| 251 | x | | | Semilandmarks running along the left side of the beak (Olsen, 2016) |
| 252 | x | x | x | Semilandmarks running along the left side of the beak (Olsen, 2016) |
| 253 | x | | | Semilandmarks running along the left side of the beak (Olsen, 2016) |
| 254 | x | x | | Semilandmarks running along the left side of the beak (Olsen, 2016) |
| 255 | x | x | x | Semilandmarks running along the left side of the beak (Olsen, 2016) |
